# A Tale of Two Positivities and the N400: Distinct neural signatures are evoked by confirmed and violated predictions at different levels of representation

**DOI:** 10.1101/404780

**Authors:** Gina R. Kuperberg, Trevor Brothers, Edward W. Wlotko

## Abstract

It has been proposed that hierarchical prediction is a fundamental computational principle underlying neurocognitive processing. Here we ask whether the brain engages distinct neurocognitive mechanisms in response to inputs that fulfill *versus* violate strong predictions at different levels of representation during language comprehension. Participants read three-sentence scenarios in which the third sentence constrained for a broad event structure, e.g. {*Agent caution animate-Patient*}. *High constraint* contexts additionally constrained for a specific event/lexical item, e.g. a two-sentence context about a beach, lifeguards and sharks constrained for the event, {*Lifeguards cautioned Swimmers*} and the specific lexical item, “swimmers”. *Low constraint* contexts did not constrain for any specific event/lexical item. We measured ERPs on critical nouns that fulfilled and/or violated each of these constraints. We found clear, dissociable effects to fulfilled semantic predictions (a reduced N400), to event/lexical prediction violations (an increased *late frontal positivity*), and to event structure/animacy prediction violations (an increased *late posterior positivity/P600*). We argue that the *late frontal positivity* reflects a large change in activity associated with successfully updating the comprehender’s current situation model with new unpredicted information. We suggest that the *late posterior positivity/P600* is triggered when the comprehender detects a conflict between the input and her model of the communicator and communicative environment. This leads to an initial failure to incorporate new unpredicted input, which may be followed by second-pass attempts to make sense of the discourse through reanalysis, repair, or reinterpretation. Together, these findings provide strong evidence that confirmed and violated predictions at different levels of representation manifest as distinct spatiotemporal neural signatures.

## Introduction

The goal of language comprehension is to extract the communicator’s intended message. This is challenging. Linguistic inputs unfold quickly, they are often ambiguous, and our communicative environments are noisy. It therefore helps if we can use context to mobilize our stored linguistic (and non-linguistic) knowledge to predict upcoming information. If incoming information matches these predictions, its processing will be facilitated. On the other hand, there will be times when we encounter inputs that violate strong predictions. In the present study, we use event-related potentials (ERPs) to ask whether the brain engages distinct neurocognitive mechanisms in response to inputs that fulfill and inputs that violate strong predictions at different levels of representation.

ERPs have provided some of the strongest evidence that the brain is sensitive to predictive processes during language comprehension. The ERP component that is primarily sensitive to *fulfilled* semantic predictions is the N400 — a centro-parietally distributed negative-going waveform that is largest between 300-500ms after the onset of an incoming word. The N400 is highly sensitive to the probability of encountering a word’s semantic features given its preceding context (Kutas & Hillyard, 1984; Kutas & Federmeier, 2011; Kuperberg, 2016).^1^ If this word’s features match semantic features that have already been predicted by a highly constraining context (e.g. as in “birthday” following context (1)), then it will evoke a smaller (less negative) N400 than a word appearing in a low constraint context (e.g. “collection” following context (2)). Importantly, however, the N400 is not a direct index of a lexical prediction violation: its amplitude is just as large to “collection” following a low constraint context, as in (2), as to “collection” when this violates a highly lexically constraining context, as in (3) (Kutas & Hillyard, 1984; Federmeier, Wlotko, De Ochoa-Dewald & Kutas, 2007).

1. He bought her a pearl necklace for her <u>birthday</u>.
2. He looked worried because he might have broken his <u>collection</u>.
3. He bought her a pearl necklace for her <u>collection</u>.

In contrast to the N400, a set of later positive-going ERP components, visible at the scalp surface between approximately 500-1000ms, do appear to be differentially sensitive to words that violate strong contextual constraints. The initial ERP research characterizing these late positivities came from different research groups working within different theoretical frameworks. One set of studies focused on a late *posteriorly* distributed positivity (maximal at parietal and occipital sites), otherwise known as the P600. This *late posterior positivity/P600* was initially characterized as a response produced by syntactic anomalies or syntactically dispreferred continuations (Hagoort, Brown, & Groothusen, 1993; Osterhout & Holcomb, 1992), but it was subsequently noted that, under certain conditions, it is also evoked by semantic incongruities — the ‘semantic P600’ (e.g. “Every morning at breakfast the eggs would *eat…”, e.g. Kuperberg, Sitnikova, Caplan & Holcomb, 2003; for a review, see Kuperberg, 2007).^2^ Another set of studies focused on a late *frontally* distributed positivity (maximal at prefrontal and frontal sites), which is typically evoked by words that violate strong lexical constraints such as in sentence (3) above (e.g. Federmeier et al., 2007; Van Petten & Luka, 2012; see also Kutas, 1993).

The studies that characterized these late positive effects used different sets of stimuli with different syntactic structures, different positions of critical words and different tasks. This made it difficult to compare the effects across different studies. However, based on a review of the early literature, Van Petten and Luka (2012) noted that, while both the late posterior and frontal positivity effects were associated with unexpected linguistic input, the main factor that distinguished them was the plausibility of the resulting interpretation. The *late posterior positivity/P600* was produced by highly implausible words. In fact, as noted by Kuperberg, 2007, section 3.4 page 32, it is typically produced by semantically *anomalous* words that result in an *impossible* interpretation (e.g. Kuperberg et al., 2003; Kuperberg et al., 2007; van de Meerendonk, Kolk, Vissers & Chwilla, 2010; Paczynski & Kuperberg, 2012). In contrast, the *late frontal positivity* is produced by unexpected but plausible critical words. This distinction was confirmed in recent studies showing that, in lexically constraining contexts, the same individuals who produced a *late frontal positivity* effect on unexpected plausible critical words, produced a *late posterior positivity* effect on unexpected highly implausible critical words (in English by DeLong, Quante and Kutas, 2014; in German by Quante, Bölte and Zwitserlood, 2018; in Hebrew by Ness & Meltzer-Asscher, 2018).

Although the research characterizing the *late posterior positivity/P600* and the *late frontal positivity* effects has proceeded somewhat independently, the mechanisms proposed to underlie each of these effects are quite similar. They include the detection of conflict between the predicted and bottom-up input (*late frontal positivity*: DeLong, Urbach, Groppe & Kutas, 2011; *late posterior positivity/P600*: van de Meerendonk, Kolk, Chwilla & Vissers, 2009; Kuperberg, 2007), the suppression of incorrectly predicted information and enhancement of activity associated with the bottom-up input (*late frontal positivity*: Federmeier et al., 2007; Kutas, 1993; *late posterior positivity/P600*: van de Meerendonk et al., 2009), and prolonged attempts to integrate the violating bottom-up input to reach a new higher-level interpretation (*late frontal positivity*: Federmeier, Kutas & Schul, 2010; DeLong, Quante & Kutas, 2014; Brothers, Swaab & Traxler, 2015; *late posterior positivity/P600*: Kuperberg, 2007; Brouwer, Fitz & Hoeks, 2012).

This leaves open many questions. For example, precisely what representations must be predicted to produce each effect? What representation must be violated by the bottom-up input to trigger each effect? And exactly why or how are such prediction violations linked to comprehenders’ interpretation of the input as plausible or impossible? The primary goal of the present study was to begin to address these questions and to bring the literatures discussing the *late frontal positivity*, the *late posterior positivity/P600*, and the N400 effects together under a common theoretical umbrella. We aimed to dissociate the two late positivity effects from one another, and from the N400 effect, by explicitly manipulating predictions that are fulfilled and violated at different grains and levels of representation in a single experiment.

In explaining the logic of our design, we appeal to three hierarchical levels of representation. At the top of the hierarchy is the comprehender’s *communication model*, which describes her beliefs about the communicator and the broader communicative environment. We assume that this communication model is of a literal English speaker transferring information about events and states that are possible in the real world (cf. Frank & Goodman, 2012; Degen, Tessler & Goodman, 2015). Within this communication model, information from the current linguistic and non-linguistic context is used to construct a *situation model* -- a high-level representation of meaning, established during deep comprehension, that describes the full set of events, actions, and characters being communicated (Van Dijk & Kintsch, 1983; Zwaan & Radvansky, 1998). Below the communication/situation model is the *event level*, which represents information about the *event structures* (sets of events that are compatible with the communication model) and the specific *events* or states that are currently being communicated. The third *semantic feature* level represents the semantic features and properties of the individual words, and sets of words, associated with these events and event structures.

We assume that these three levels are distinguished in at least two ways. The first is by the *time scale* of the linguistic input with which they interact: in general, it requires more time (and linguistic information) to build a rich situation model than to infer a single event; similarly, it requires more time/linguistic information to infer a whole event than the meaning of a single word. Because of this, information that is fully encoded at a higher level of the hierarchy will subsume information encoded at a lower level. Second, we assume that information represented at each of these different levels is encoded in a different form, and therefore that it interacts with different *types* of linguistic information. For example, the semantic features level can receive input from phonological or orthographic inputs that are used to decode the semantic features associated with specific words, while the event level can interact with sets of semantic features (e.g. animacy information) and certain syntactic cues, which provide information about an event structure, as well as with finer-grained features that provide information about a specific event or state.

We assume that, during language comprehension, as new linguistic information becomes available and incrementally decoded, it is passed up the hierarchy in a bottom-up fashion. In addition, information can flow down the hierarchy in a top-down fashion. Specifically, the communication model provides top-down constraints determining which events are possible at the event level (the full set of possible event structures), while the situation model influences the probability/degree of activation over specific events. Similarly, the probability/degree of activation of specific events can influence the probability/degree of activation over semantic features represented at the lower semantic features level. Within this dynamic framework, if information from a higher level reaches and changes the state of information represented at a lower level before new bottom-up linguistic information arrives and is decoded at that lower level, we refer to this as top-down *prediction*. In other words, we use the term, *prediction,* both in a spatial sense (top-down effects from a higher to a lower level of representation) and in a temporal sense (input at time point *t* changes the state of activity at *t*+1).

We conceptualize the N400 as reflecting access to the semantic features associated with new bottom-up inputs that have not already been predicted — that is, as changes in activity at the semantic feature level that are induced by new inputs, with smaller changes associated with inputs whose semantic features have already been predicted by the prior context (see Kuperberg, 2016). We hypothesize that the *late frontal positivity* reflects a large change in activity associated with successfully updating the comprehender’s current situation model, and that this component is triggered when new bottom-up input violates prior top-down predictions that originated from the prior situation model. By updating the situation model, the source of these earlier incorrect predictions can be corrected. We suggest that the *late posterior positivity/P600* is evoked when new bottom-up input *conflicts* with the constraints of the communication model itself, which prevents the input from being initially incorporated into the current situation model. This conflict and resulting initial interpretive failure may lead to additional second-pass attempts to make sense of the input (see Shetreet, Alexander, Romoli, Chierchia & Kuperberg, 2019 for recent discussion).

### Design of the current study

As a step towards testing this theory, we created a set of well-controlled three-sentence discourse scenarios. These contexts were written so that, just before the onset of a critical word, they constrained for different types of information at the levels of representation described above. This is illustrated schematically in Figure 1.

1A. *Low constraint context:* Eric and Grant received the news late in the day. They mulled over the information, and decided it was better to act sooner rather than later. Hence, they cautioned the…(trainees/drawer).
1B. *High constraint context:* The lifeguards received a report of sharks right near the beach. Their immediate concern was to prevent any incidents in the sea. Hence, they cautioned the…(swimmers/trainees/drawer).

**Figure 1.**
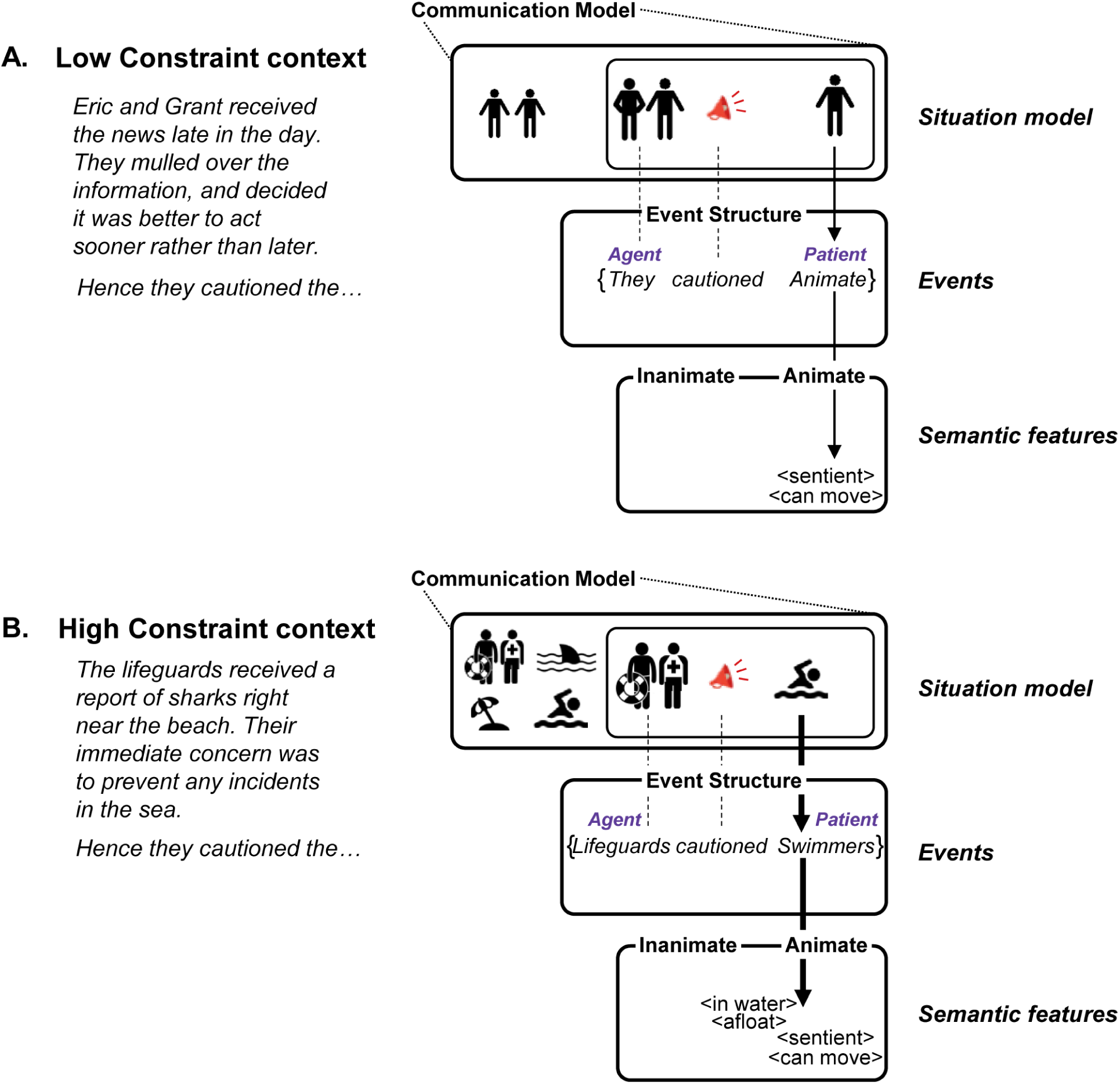
Schematic illustration of the state of the language comprehension system at three hierarchical levels of representation after reading a *low constraint* context and a *high constraint* context, but before encountering the upcoming critical noun. Please see text pages 10-11 for full explanation. Icons used in this and Figures 7 and 8 were obtained from thenounproject.com

*Low constraint* contexts established a relatively simple situation model. For example, in 1A, the first two sentences simply introduce two unknown characters, Eric and Grant, who are considering taking an unknown action (see Figure 1A, left side of situation model). The first few words of the third sentence (“They cautioned the…”) establishes that Eric and Grant are about to caution an unknown person/people (see Figure 1A, right side of situation model). While this situation model constrains strongly for a particular event structure, {*Agent cautioned animate-Patient*}, it does not constrain for any specific event.^3^ The {*Agent cautioned animate-Patient*} event structure, in turn, constrains for semantic features that are typically associated with animate entities (e.g. <sentient>, <can move>).

*High constraint* contexts established a rich situation model. For example, in 1B, the first two sentences establish the presence of lifeguards and sharks in a beach scene, which is likely to include swimmers (see Figure 1B, left side of situation model). This means that the same first few words of the third sentence (“They cautioned the…”) establish that the lifeguards are highly likely to caution a group of swimmers (Figure 1B, right side of situation model). Thus, in these *high constraint* contexts, the situation model constrains not only for a particular event structure, {*Agent cautioned animate-Patient*}, but also for a specific event, {*Lifeguards cautioned Swimmers*}. This specific event, in turn, constrains not only for semantic features that are typically associated with animate entities (e.g. <sentient>, <can move>), but also for features that are more specifically associated with the lexical item, *swimmers* (e.g. <in water>, <afloat>).

Following each context, we introduced critical words in the direct object position of the third sentence, which either confirmed or violated each of these constraints. This yielded five types of discourse scenarios (see Table 1 for examples). In the *high constraint expected* scenarios, the critical word (“swimmers”) was highly predictable because it satisfied all top-down constraints at both the event level and at the semantic features level. In all four other conditions, the critical words were unpredictable, but each for a different reason. In the *low constraint unexpected* scenarios, the critical word (“trainees”) was plausible but unexpected because the discourse context did not constrain for this item, or any other single event or lexical item. The critical word also did not violate event structure/animacy constraints of the prior verb. In the *high constraint unexpected* scenarios, the critical word (“trainees”) was plausible but unexpected because it violated constraints for a specific event, {*Lifeguard cautioned Swimmers*}, and for the specific semantic features associated with the expected word (“swimmers”). Again, it did not violate event structure/animacy constraints. In the *low constraint anomalous* scenarios, the critical word (“drawer”) was anomalous because it violated event structure/animacy constraints — its inanimate features were incompatible with an animate Patient. However, it did not violate constraints for a specific event or for the semantic features associated with a specific lexical item because the discourse context did not constrain strongly for any single event/lexical item. Finally, in the *high constraint anomalous* scenarios, the critical word (“drawer”) produced a “double violation”, violating *both* event structure/animacy constraints as well as constraints for a specific event/lexical item.

**Table 1.** Examples of five experimental conditions created around the same verb (here, “cautioned”). The critical word in each of the example final sentences is underlined (although this was not the case in the experiment itself). The final sentences continued with additional words, as indicated by the three dots. Means are shown with standard deviations in parentheses. *SSV: Semantic Similarity Value: the cosine similarity between vector corresponding to the critical word and the vector corresponding to the preceding context, extracted using Latent Semantic Analysis using the pairwise comparison tool (term-to-document) at http://lsa.colorado.edu/ (possible range of SSVs: +1 to −1).

| Scenario Type | Example | Lexical constraint of context | Cloze of critical word | SSV* between critical word and preceding context |
| --- | --- | --- | --- | --- |
| **1. High Constraint Expected** | The lifeguards received a report of sharks right near the beach. Their immediate concern was to prevent any incidents in the sea. Hence, they cautioned the <u>swimmers</u>... | 71%<br>(13%) | 71%<br>(13%) | 0.18<br>(0.18) |
| **2. Low Constraint Unexpected** | Eric and Grant received the news late in the day. They mulled over the information, and decided it was better to act sooner rather than later. Hence, they cautioned the <u>trainees</u>... | 20%<br>(9%) | 0.2%<br>(0.8%) | 0.01<br>(0.05) |
| **3. High Constraint Unexpected** | The lifeguards received a report of sharks right near the beach. Their immediate concern was to prevent any incidents in the sea. Hence, they cautioned the <u>trainees</u>... | 71%<br>(13%) | 0.1%<br>(0.5%) | 0.01<br>(0.05) |
| **4. Low Constraint Anomalous** | Eric and Grant received the news late in the day. They mulled over the information, and decided it was better to act sooner rather than later. Hence, they cautioned the <u>drawer</u>... | 20%<br>(9%) | 0%<br>(0%) | 0.01<br>(0.06) |
| **5. High Constraint Anomalous** | The lifeguards received a report of sharks right near the beach. Their immediate concern was to prevent any incidents in the sea. Hence, they cautioned the <u>drawer</u>... | 71%<br>(13%) | 0%<br>(0%) | 0.01<br>(0.04) |

Critical words in the four unpredictable scenarios were matched on their general semantic relatedness with the preceding content words in their contexts, as assessed using Latent Semantic Analysis (LSA: Landauer & Dumais, 1997; Landauer, Foltz, & Laham, 1998). Participants read and monitored the coherence of the scenarios, and we measured ERPs on the critical words.

Our first set of questions concerned the N400. Based on numerous previous studies, we expected that the N400 would be smaller on critical words in the *high constraint expected* scenarios than in all four types of unpredictable scenarios. Based on the work by Kutas & Hillyard (1984) and Federmeier, Wlotko, De Ochoa-Dewald & Kutas (2007), we also predicted that, in the plausible scenarios, the N400 evoked by unpredictable critical words would be insensitive to the lexical constraint of the context (no difference between the *high constraint unexpected* and the *low constraint unexpected* conditions). In addition, our design allowed us to ask whether the amplitude of the N400 evoked by semantically anomalous critical words would also be insensitive to the lexical constraint of the context (predicting no difference between the *high constraint anomalous* and the *low constraint anomalous* conditions). Finally, we were interested in whether the N400 would be larger on the anomalous than the unexpected but plausible critical words. If so, this would provide evidence that the N400 evoked by a given word can be sensitive to its implausibility (here, operationalized by whether that word matched or mismatched the verb’s animacy selection restrictions), even when its cloze probability and its semantic relatedness with its preceding content words are matched across conditions (see Paczynski & Kuperberg, 2011, 2012).

Our second set of questions concerned the late positivities. We hypothesized that we would be able to dissociate the scalp distribution of these positivities based on whether they violated strong predictions for a single event/lexical item or for a whole event structure. Specifically, we hypothesized that the *high constraint unexpected* critical words, which violated strong constraints for a specific event/lexical item, would selectively evoke a *late frontal positivity* (larger than that produced by critical words in any other condition), while the *low constraint anomalous* critical words, which violated strong constraints for an event structure/animacy features, would selectively evoke a *late posterior positivity/P600* (larger than that produced by critical words in any of the plausible scenarios). This would provide strong evidence that distinct neurocognitive mechanisms can be triggered by violations at these different grains of representation.

Finally, our design allowed us to ask not only whether the neurocognitive mechanisms underlying the *late frontal positivity* and the *late posterior positivity/P600* are distinct, but also whether they are independent of one another. This would, in turn, constrain our understanding of the precise levels of representation at which violations must be detected to trigger each of these effects (see Discussion). The key condition to address this question was the doubly violated scenarios (*high constraint anomalous*) in which the critical words violated *both* constraints for a specific event/lexical item as well as for a broader event structure. One possibility is that the neurocognitive processes underlying the *late frontal positivity* and the *late posterior positivity/P600* are independent of one another. For example, if the *late frontal positivity* only reflects the detection of a violated prediction at the level of semantic features, while the *late posterior positivity/P600* reflects the detection of a violated event structure, then, assuming additivity of independent ERP effects (see Osterhout & Nicol, 1999), one might expect the doubly violated critical words to evoke *both* a *late frontal positivity* and a *late posterior positivity/P600*. If, on the other hand, the neurocognitive processes underlying the *late frontal positivity* and the *late posterior positivity/P600* are interdependent, then we should see non-additive effects. For example, if the detection of an event structure violation interferes with processes reflected by the *late frontal positivity*, then we should see a *late posterior positivity/P600* but no *late frontal positivity* on the double violations.

## Methods

### Development and norming of materials

Participants read the five types of scenarios discussed above (see Table 1). In all scenarios, the first two sentences introduced a context. The third sentence began with an adjunct phrase of 1-4 words, followed by a pronominal subject that referred back to the first two sentences, followed by a verb, a determiner, a *critical word* (always a direct object noun), and then three additional words. The scenarios varied by the constraint of the context (i.e., the combination of the first two sentences and the third sentence until just before the critical word), and by whether the critical word matched or violated the event/lexico-semantic constraints of the high constraint contexts and/or the event structure/animacy constraints defined by the thematic-semantic properties of the previous verb.

To construct these scenarios, we began with a set of 100 preferentially transitive verbs (50% selecting for animate direct objects; 50% for inanimate direct objects), which, in the absence of a discourse context, were not highly predictive of any particular upcoming event or noun. The constraints of these verbs in minimal context were assessed in an offline cloze norming study (see below). We then wrote high and low constraint two-sentence discourse contexts around each verb (matching the average number of words across the two levels of constraint), and carried out a second cloze norming study of these discourse contexts (see below). Based on the results of this cloze norming, we created the five scenario types. To create the *high constraint expected* scenarios, each *high constraint* context was paired with the noun with the highest cloze probability for that context. To create the *high constraint unexpected* scenarios, each *high constraint* context was paired with a direct object noun of zero (or very low) cloze probability, but that was still plausible in relation to this context. To create the *low constraint unexpected* scenarios, the same unexpected noun was paired with the *low constraint* context. To create the *anomalous* scenarios, each *high constraint* and *low constraint* context was paired with a noun that violated the animacy selectional restrictions of the verb. Thus, one of the five conditions had predictable critical words, and four had unpredictable critical words.

In constructing these scenarios, we used Latent Semantic Analysis (LSA, Landauer & Dumais 1997; Landauer et al. 1998) to match the semantic similarity between the critical words and entirety of the prior contexts across the four unpredictable conditions. To carry out this analysis, we used the pairwise comparison tool from http://lsa.colorado.edu/ with default values (local and global weighting functions, 300 latent semantic dimensions). We extracted pair-wise *term-to-document* Semantic Similarity Values (SSVs) — cosine similarities between the vectors corresponding to our critical words and ‘pseudo-document’ vectors corresponding to each of our contexts. All of our critical words appeared in the original term-document matrix, except for five for which we substituted close synonyms. The mean SSVs for each of the five types of scenario are shown in Table 1. SSVs showed no significant difference across the four unpredictable conditions, *F*(3,396) = 0.173, *p* = .914. Unsurprisingly, the SSVs were significantly greater in the predictable scenarios than in each of the four unpredictable scenarios (all pair-wise comparisons, *p*s < .001, *t*s > 9).

#### Cloze norming studies

As noted above, we carried out two cloze norming studies to construct the stimuli. For both studies, participants were recruited through the crowd-sourcing platform, Amazon Mechanical Turk. They were asked to complete each context with the first word that came to mind (Taylor, 1953), and, in an extension of the standard cloze procedure, to also provide two additional words that could complete the sentence (see Federmeier et al, 2007; Schwanenflugel & LaCount, 1988). Responses were excluded from participants who reported that their first language learned was not English or if they reported any psychiatric or neurological disorders. Responses were also excluded from any participants who failed to follow instructions (“catch” questions were used as periodic attention checks).

##### Cloze norming study 1: Selection of non-constraining verbs

We started with a set of 600 transitively-biased verbs, compiled from various sources including Levin (1993) and materials from previous studies conducted in our laboratory (Paczynski & Kuperberg, 2011, 2012). Verbs with log Hyperspace Analogue to Language (HAL) frequency (Lund & Burgess, 1996) of two standard deviations below the mean (based on English Lexicon Project database, Balota et al. 2007) were excluded. For each verb, we constructed a simple active, past tense sentence stem that consisted of only a proper name, the verb, and a determiner (e.g., “Harry explored the…”). These sentences were divided into six lists in order to decrease the time demands on any individual participant during cloze norming. Between 89 and 106 participants (depending on list) who met inclusionary criteria provided completions for each verb.

The lexical constraint of each verb in these minimal contexts was calculated by identifying the most common completion across participants and tallying the proportion of participants who provided this completion. The set of verbs used in the final set of stimuli had an average constraint of 16.4% (*SD* = 0.7%).

##### Cloze norming study 2: Selection and characterization of the final set of discourse stimuli

We began with an initial set of 198 pairs of *high constraint* and *low constraint* contexts (the combination of the first two sentences and the third sentence, including the verb and the determiner). These were pseudorandomly divided into two lists, such that each list contained only one of the two contexts associated with each verb. The two lists were then divided into thirds to decrease time demands on any individual participant during cloze norming. Between 51 and 69 participants who met inclusionary criteria provided completions for each scenario.

Cloze probabilities were calculated based on the percentage of respondents providing the critical noun used in the experiment. Alternate word forms (e.g., singular/plural) were collapsed, but synonyms or lexical alternatives were not collapsed (e.g. couch/sofa). The lexical constraints of the full discourse contexts were calculated as described above. The mean and standard deviation of the cloze probability and the lexical constraint for each scenario type is given in Table 1.

#### Counterbalancing across lists

The final set of 500 scenarios (100 sets of five) were rotated across the five lists, with 20 items per condition in each list. Counterbalancing ensured that the same adjunct phrase plus verb in each of the final sentences appeared in all conditions across the five lists, and the same critical words were counterbalanced across the four types of unpredictable scenarios (*low constraint unexpected, high constraint unexpected, low constraint anomalous, high constraint anomalous*). The critical words in the *high constraint expected* scenarios differed from those in the four types of unpredictable scenarios (see Supplementary Materials, Table S1 for information about the lexical characteristics of these critical words). The same *high constraint* contexts were counterbalanced across the *high constraint expected*, *high constraint unexpected* and *high constraint anomalous* scenarios, and the same *low constraint* contexts were counterbalanced across the *low constraint unexpected* and *low constraint anomalous* scenarios.

To each list, we then added an additional 20 *high constraint anomalous*, 20 *low constraint unexpected*, and 20 *low constraint anomalous* filler scenarios so that each participant saw 160 scenarios in total. This ensured that each participant saw an equal number of high and low constraint contexts, and that, at each level of contextual constraint (*high constraint* versus *low constraint*), half of the scenarios were plausible and the other half were anomalous.

### Participants

We report data from thirty-nine native English speakers (age 18-32 years, mean: 21.6, *SD*: 3.6; 21 male). Forty participants were originally recruited but one failed to complete the session. Participants were recruited from Tufts University and the surrounding communities. They were screened on the basis of the following exclusion criteria: significant exposure to any language other than English before the age of five years; history of psychiatric or neurological diagnoses or injury; use of psychoactive medication within the preceding 6 months. Participants were right-handed, as assessed by the Edinburgh Handedness Inventory (Oldfield, 1971). They provided written informed consent, were paid for their time, and all protocols were approved by Tufts University Social, Behavioral, and Educational Research Institutional Review Board.

### Stimulus presentation and Task

Participants sat in a dimly lit room while stimuli were presented approximately 150cm from the computer screen. Stimuli were presented using PsychoPy 1.83 software (Peirce, 2007), and were displayed on a LCD monitor using white Arial font set to 0.1 of the screen height, on a black background. Details of the structure of each trial are given in Figure 2 and legend.

**Figure 2.**
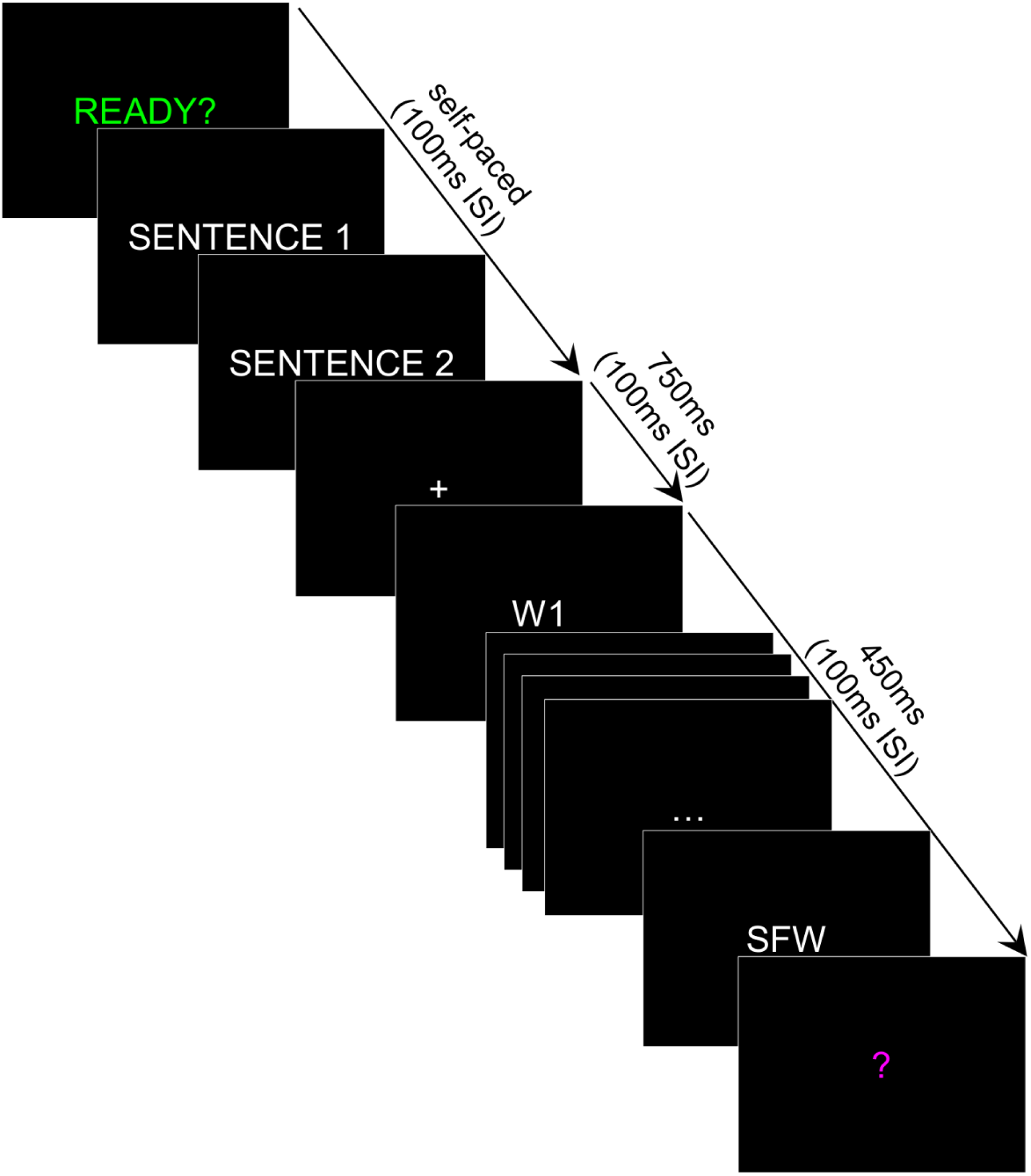
Presentation of stimuli. Each trial began with the prompt “READY?” (centered in green font). When participants initiated the trial with a button press, the first two sentences were presented successively in full on the screen. Participants advanced through these sentences at their own pace using a button press. Then a centered fixation cross appeared (750ms; 100ms inter-stimulus interval, ISI), followed by the third sentence, which was presented word-by-word (each word 450ms; 100ms ISI). After the sentence-final word (SFW), a “?” appeared (centered, magenta font), prompting participants to make an acceptability judgement about whether or not the preceding scenario “made sense” or not. The “?” remained on the screen until the response was registered.

Participants’ task was to press one of two buttons after seeing a ‘‘?’’ cue to indicate whether they judged each scenario to “make sense” or not. This task encouraged active coherence monitoring during online comprehension and was intended to prevent participants from completely disregarding the anomalies (see Sanford et al. 2011 for evidence that detecting anomalies is necessary to produce a *late positivity/P600* effect at all, and see https://projects.iq.harvard.edu/kuperberglab/additional-materials/P600-task-discussion, for a discussion of the use of this task in ERP studies). In addition, following approximately 32 trials of the 160 trials (all fillers), participants answered a yes/no comprehension question about the preceding scenario. For example, the scenario, “Damien was not at all like himself that day. Something in the atmosphere made him act differently, and everyone could tell. Relentlessly, he ridiculed the patients until they cried.” Was followed by the question, “Is this behavior typical of Damien?”. This encouraged participants to attend to the scenarios as a whole rather than just the third sentence in which the anomalies appeared. The experiment was divided into four blocks, with block order randomized across participants. Participants were given ten practice trials before the main experiment.

### EEG recording and processing

EEG was recorded using a Biosemi Active-Two acquisition system from 32 active electrodes in a modified 10/20 system montage. Signals were digitized at 512 Hz and a passband of DC - 104 Hz. All processing was carried out using EEGLab (Delorme & Makeig, 2004) and ERPLab (Lopez-Calderon & Luck, 2014) analysis packages in the MATLAB environment. After importing the data for processing, the EEG was referenced offline to the average of the left and right mastoid channels. Both a high-pass (0.05Hz) and a low-pass (30Hz) 24dB per octave filter were applied offline to the continuous data.

The EEG was then segmented into initial epochs spanning from −300ms before until +1400ms after the presentation of critical words. Independent Components Analysis (using the extended infomax algorithm implemented in EEGLab) was then applied to the epoched data to correct for blinks and other eye movement artifact. Then, trials with any residual artifacts were rejected using a semi-automated procedure with participant-specific artifact detection thresholds. After artifact rejection, there remained on average 18.6 trials per condition (*SD* = 1.3). Artifact rejection rates did not differ significantly across the five conditions, *F*(4,152) = 1.91, *p* = .13. Our epoch of interest was −100-1000ms.

We then extracted single trial artifact-free ERP data, using a baseline of −100-0ms, by averaging across electrode sites and time windows that defined specific spatiotemporal regions of interest that we selected *a priori* to operationalize each of our three ERP components of interest, see Figure 3. Based on numerous previous studies, we operationalized the N400 as the average voltage within the central region between 300-500ms. In the prior literature, the two late positivities have shown variable time courses (e.g. Federmeier et al., 2007; DeLong, Quante and Kutas, 2014; Kuperberg et al., 2003;Paczynski & Kuperberg, 2011, 2012), and so, conservatively, we operationalized both of these components as the average voltage across a long 600-1000ms time window. For the *late frontal positivity*, we averaged across electrode sites within the prefrontal region (based on Federmeier et al. 2007), and for the *late posterior positivity/P600*, we averaged across electrode sites within the posterior region (based on Kuperberg et al., 2003; Paczynski & Kuperberg, 2011, 2012).

**Figure 3.**
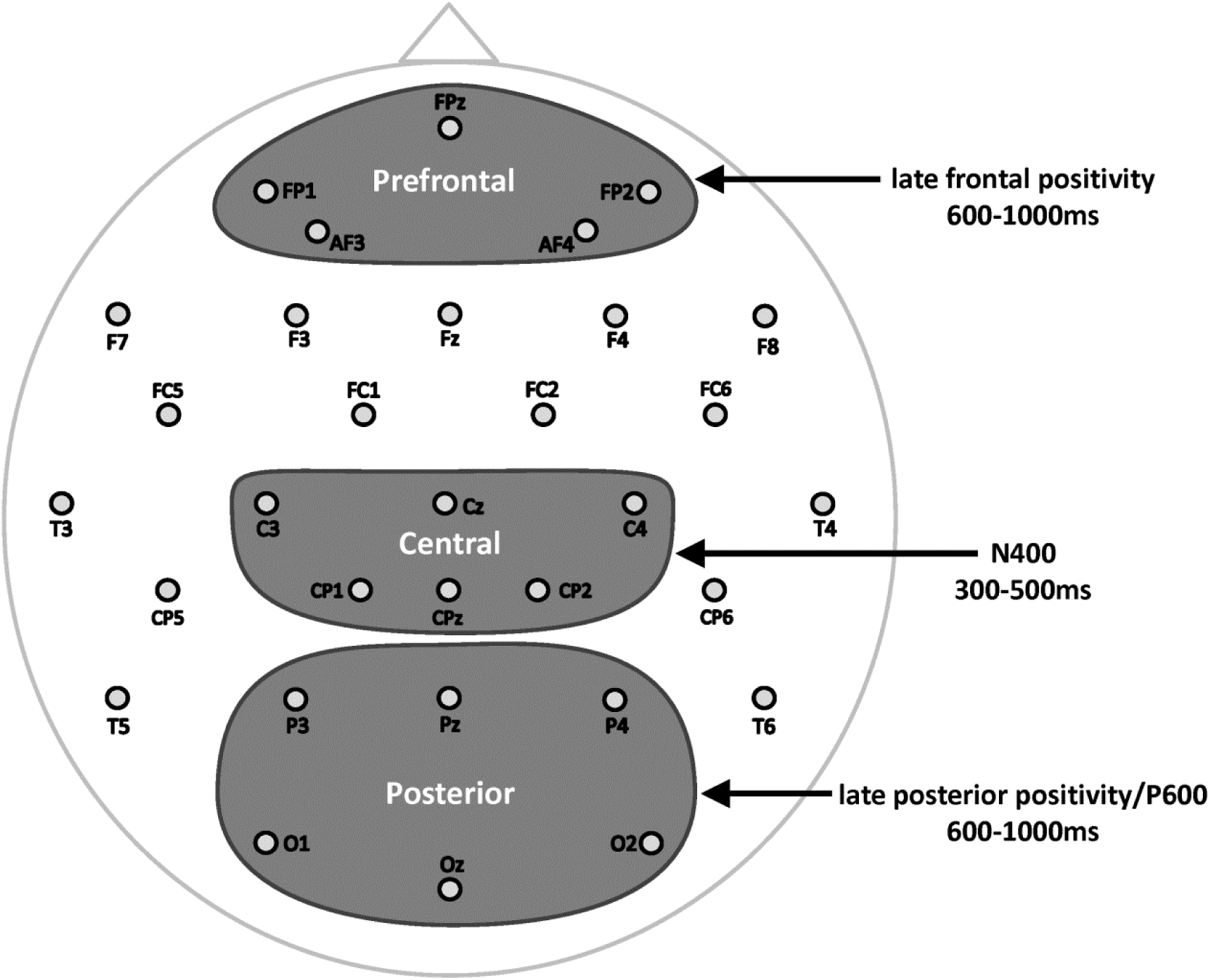
Spatiotemporal regions of interest used for analysis. The N400 was operationalized as the average voltage between 300-500ms across all electrode sites within the central region. The *late frontal positivity* was operationalized as the average voltage (600-1000ms) in the prefrontal region. The *late posterior positivity/P600* was operationalized as the average voltage (600-1000ms) in the posterior region

### ERP statistical analysis

We report the results of a series of linear mixed-effect regression models, fit in R version 3.2.4 (R Core Team, 2016) using the “lme4” package version 1.1-11 (see Bates et al., 2015). Our dependent measure was the average trial-level ERP amplitude within each of the spatiotemporal regions of interest described above.^4^ The maximal random effects structure was used (Barr, Levy, Scheepers, & Tily, 2013) with by-item and by-subject random intercepts, and by-item and by-subject random slopes for each fixed effect of interest. For each contrast of interest, *p*-values were estimated using a Satterthwaite approximation, as implemented by the “lmerTest” package version 2.0-30 (Kuznetsova et al., 2015).

On the N400, we were primarily interested in whether, across the four unpredictable conditions, there were main effects of Constraint, Plausibility, and/or an interaction between these two factors, and so we constructed a model that crossed two levels of Constraint (*high constraint*, *low constraint*) and two levels of Plausibility (*plausible*, *anomalous*). We also carried out pair-wise comparisons between each of the four unpredictable conditions and the *high constraint expected* condition to confirm the presence of significant N400 effects.

On the late positivities, we were primarily interested in the contrasts between specific unpredictable conditions and all other conditions. We therefore proceeded straight to planned pair-wise comparisons. For the *late frontal positivity*, we were interested in the comparison between the *high constraint unexpected* critical words and each of the four other conditions, and for the *late posterior positivity/P600*, we were interested in the comparison between the anomalous critical words (both *high constraint anomalous* and *low constraint anomalous*) and each of the three plausible conditions. For both late positivities, we carried out additional pairwise comparisons between all remaining conditions to determine the specificity of any effects.

## Results

### Behavioral findings

On average, participants correctly judged the acceptability of the prior scenario on 89.6% of trials. Participants were most accurate in responding “YES” to the *high constraint expected* scenarios (97.6%), less accurate in responding “NO” to the *high constraint anomalous* (91.0%) and the *low constraint anomalous* (89.8%) scenarios, and least accurate in responding “YES” to the *high constraint unexpected* and the *low constraint unexpected* scenarios (both 86.3%). Participants correctly answered 86.4% of the comprehension questions on average (27.6 out of 32 questions; *SD*: 2.0), indicating that they attended to the full contexts of the scenarios, rather than just to the final sentences.

### ERP findings

#### N400

Figure 4 shows grand-average ERP waveforms at electrode site, Cz, for each of the five conditions. It also shows the mean voltage (averaged across the central 300-500ms spatiotemporal region used to operationalize the N400) for each condition. As expected, the N400 was smaller (less negative) on critical words in the *high constraint expected* scenarios than on critical words in each of the four types of unpredictable scenario (all *t*s > 6.0, all *p*s < .001).

**Figure 4.**
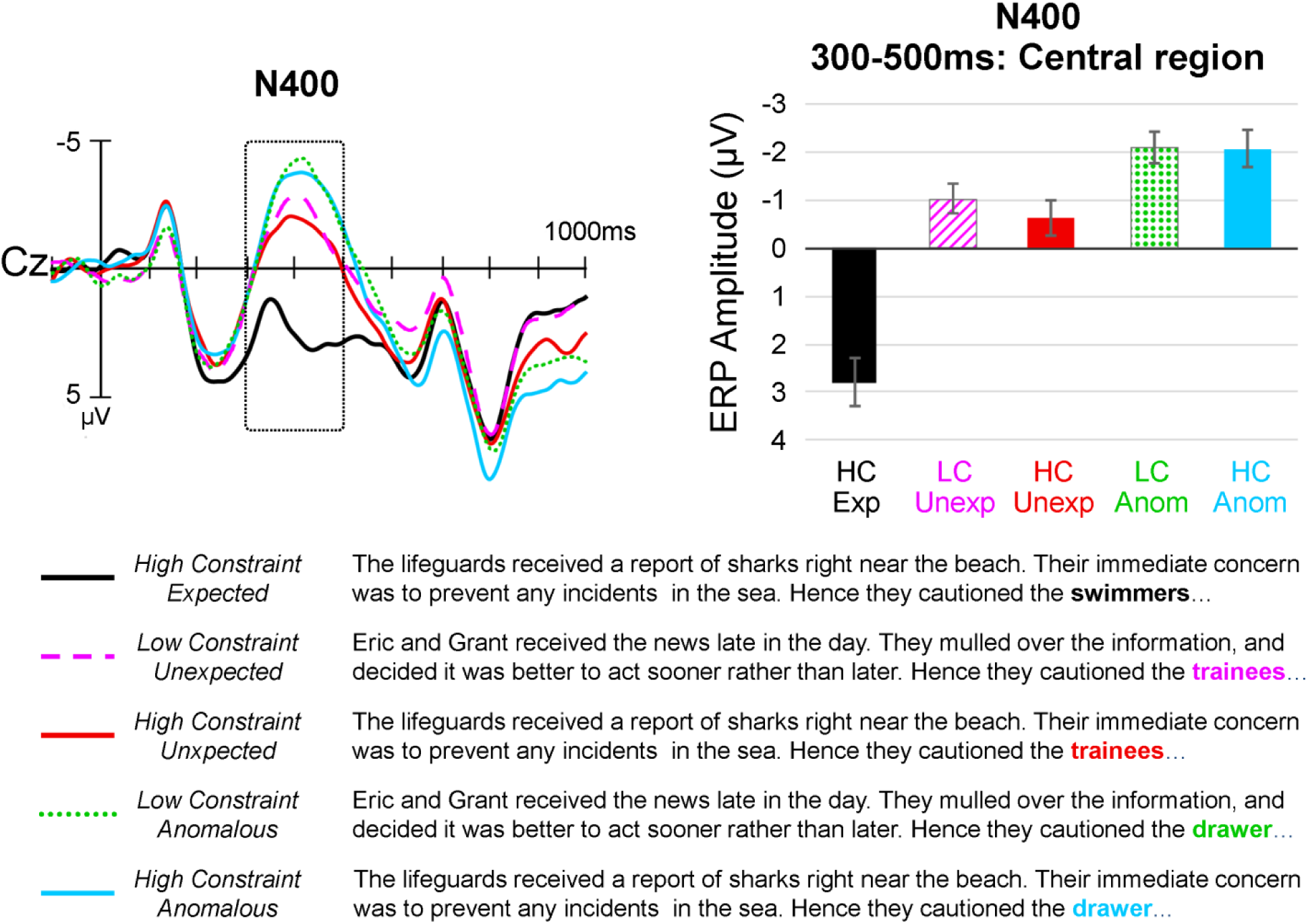
N400. Left: Grand-averaged ERP waveforms at electrode Cz in each of the five conditions. Right: Mean N400 amplitude in each of the five conditions, with the N400 operationalized as the average voltage across all time points between 300-500ms across all electrode sites within the central region of interest. Negative voltage is plotted upward. Error bars represent ±1standard error of the mean, calculated within-subjects (Morey, 2008).

Across the four types of unpredictable scenarios, a model that crossed Constraint (*high constraint*, *low constraint*) and Plausibility (*plausible*, *anomalous*) showed no main effect of Constraint (*t* = −0.54, *p* = .59), no interaction between Constraint and Plausibility (*t* = 0.31, *p* = .75), but a significant main effect of Plausibility (*t* = −3.08, *p* = .003) due to a slightly larger (more negative) N400 on the anomalous than the plausible critical words. Voltage maps for all pair-wise comparisons are shown in Supplementary Materials, Figure S1, and the full set of statistical comparisons between all pairs of conditions is reported in Supplementary Materials, Table S2.

#### Late frontal positivity

Figure 5 (top) shows grand-average ERP waveforms at electrode site FPz in each of the five conditions. It also shows the mean voltage (averaged across the prefrontal 600-1000ms spatiotemporal region used to operationalize the *late frontal positivity*) for each condition. As hypothesized, the *late frontal positivity* was larger (more positive) to critical words that violated strong event/lexical constraints (*high constraint unexpected* scenarios) than to critical words in all other conditions (all *t*s > 2.3, all *p*s < .02). This effect was selective: pair-wise contrasts that did not include the *high constraint unexpected* condition failed to show any significant effects on the *late frontal positivity* (all *t*s < 1.7, all *p*s > .11). Of particular note, no significant *late frontal positivity* effect was produced by the critical words in the *high constraint anomalous* scenarios, which violated *both* event/lexical and event structure/animacy constraints, relative to the *low constraint anomalous*, the *high constraint expected* or the *low constraint unexpected* critical words (all *t*s < 1.0, *p*s > .49). Voltage maps that illustrate the scalp distribution of the *late frontal positivity* effects are shown in Figure 6A. The full set of statistical comparisons between all pairs of conditions is reported in Supplementary Materials, Table S3.

**Figure 5.**
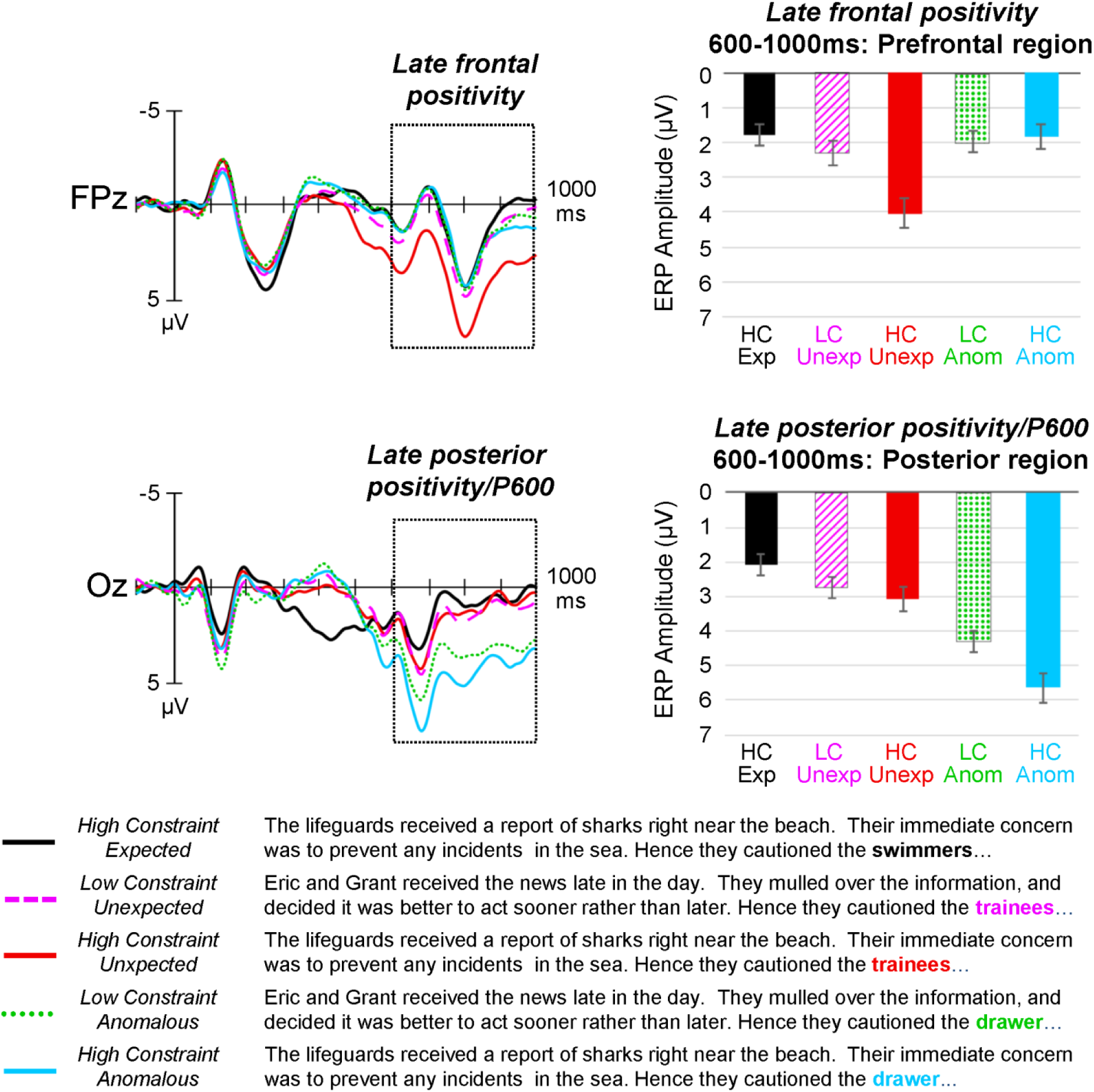
The late positivities. Top: The *late frontal positivity.* Top left: Grand-averaged ERP waveforms at electrode FPz in each of the five conditions. Top right: Mean *late frontal positivity* amplitude in each of the five conditions, with the *late frontal positivity* operationalized as the average voltage across all time points between 600-1000ms across all electrode sites within the prefrontal region of interest. Bottom: The *late posterior positivity/P600.* Bottom left: Grand-averaged ERP waveforms at electrode Oz in each of the five conditions. Bottom right: Mean *late posterior positivity/P600* amplitude in each of the five conditions, with the *late posterior positivity/P600* operationalized as the average voltage across all time points between 600-1000ms across all electrode sites within the posterior region of interest. Negative voltage is plotted upward. Error bars represent ±1standard error of the mean, calculated within-subjects (Morey, 2008).

**Figure 6.**
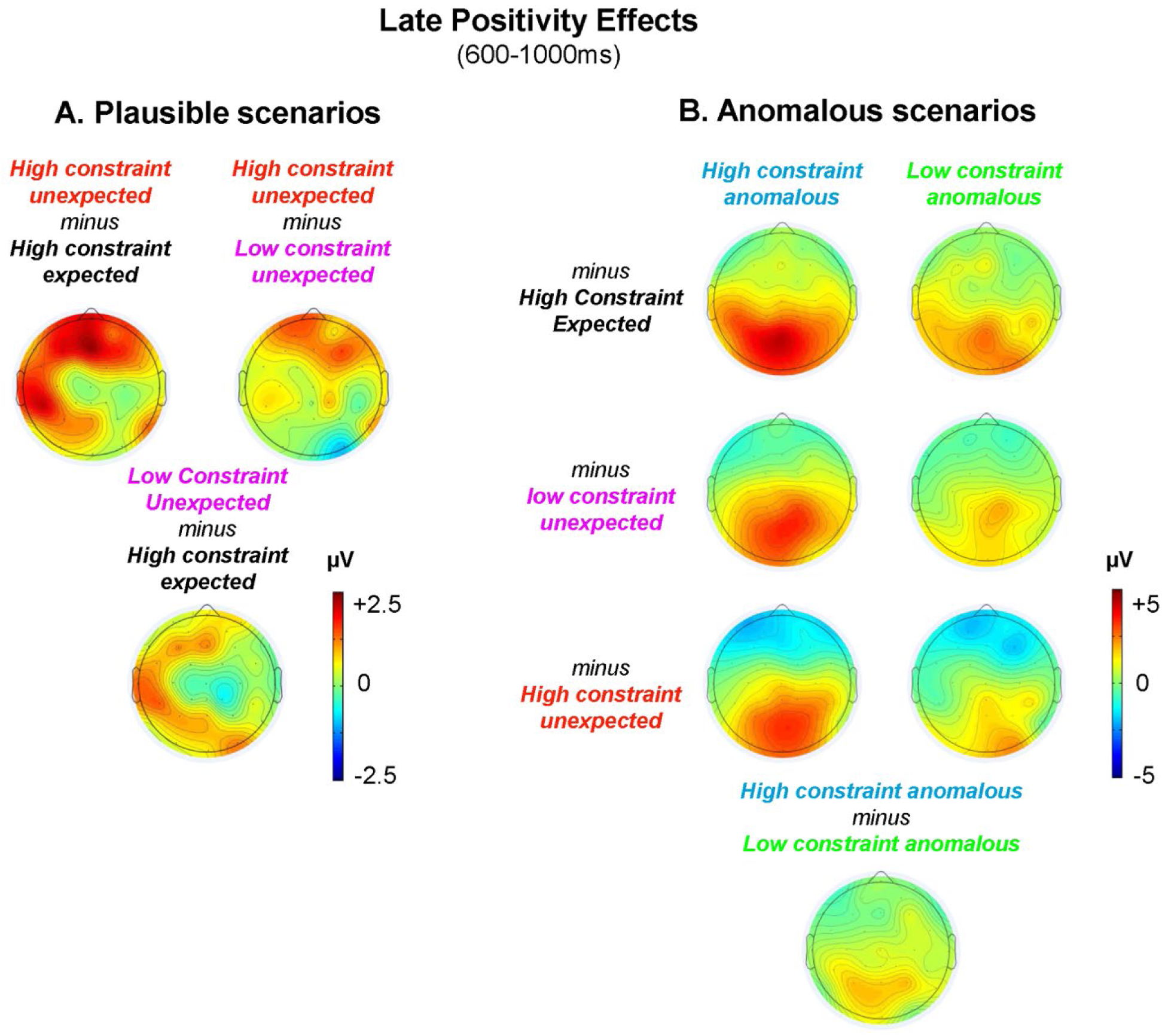
Voltage maps showing the scalp topographies of the late positivity effects. Voltages are averaged across the 600-1000ms time window for all pair-wise contrasts. **A.** *Left:* Voltage maps contrasting each of the plausible conditions with one another. These illustrate the frontal scalp distribution of the late positivity effect produced by the *high constraint unexpected* critical words, relative to the *high constraint expected* critical words, and relative to the *low constraint unexpected* critical words. The scalp distribution of the effect produced by the *low constraint unexpected*, relative to the *high constraint expected* critical words, is similar, but the effect was not statistically significant. **B.** *Right:* Voltage maps contrasting the anomalous critical words with critical words in each of the plausible conditions. These illustrate the posterior scalp distribution of the late positivity effect evoked by the anomalous critical words. The effects produced by the *high constraint anomalous* critical words were larger than those produced by the *low constraint anomalous* critical words. Note that the *late frontal positivity* effects (left) and the *late posterior positivity/P600* effects (right) are shown at different voltage scales to better illustrate the scalp distribution of each effect.

#### Late posterior positivity/P600

Figure 5 (bottom) shows grand-average ERP waveforms at electrode site Oz in each of the five conditions. This figure also shows the mean voltage (averaged across the posterior 600-1000ms spatiotemporal region used to operationalize the *late posterior positivity/P600*) for each condition. As hypothesized, the *late posterior positivity/P600* was larger (more positive) to anomalous critical words that violated event structure/animacy constraints (both *high constraint anomalous* and *low constraint anomalous*) than to plausible critical words (*high constraint expected, low constraint unexpected* and *high constraint unexpected* critical words), all *t*s > 3.7, all *p*s < .001. Again, this effect was selective: pair-wise comparisons that did not include the anomalous conditions failed to reveal any significant effects on the *late posterior positivity/P600* (all *t*s < 1.83, all *p*s > .07). Of particular note, the *high constraint anomalous* scenarios, which violated *both* event/lexical and event structure/animacy constraints, produced a larger *late posterior positivity/P600* than the *low constraint anomalous* scenarios, which violated just event structure/animacy constraints (*t* = 2.31, *p* = .03). Voltage maps that illustrate the scalp distribution of the *late posterior positivity/P600* effects are shown in Figure 6B. The full set of statistical comparisons between all pairs of conditions is reported in Supplementary Materials, Table S4.

## Discussion

We used a single set of well-controlled stimuli to examine neural activity in response to words that confirmed and violated constraints at different levels of representation during language comprehension. Our findings were clear. Words that confirmed predictions of semantic features produced an attenuated N400; words that violated event/lexical constraints produced an enhanced *late frontal positivity*, while words that that violated event structure/animacy constraints elicited an enhanced *late posterior positivity/P600*. These findings suggest that the brain recognized and distinguished the different levels and grains of representation that were predicted and violated as a consequence of our experimental manipulations. In the sections below, we discuss these findings in more detail, and how they can be accommodated within a *hierarchical generative framework* of language comprehension (see Kuperberg & Jaeger, 2016; Kuperberg, 2016; see also Xiang & Kuperberg, 2015). We then return to the functional significance of each of these ERP components, and discuss the broader theoretical implications of our findings.

### The N400: Confirmed predictions at the level of semantic features

As expected, the N400 was significantly smaller to semantically predictable critical words in the *high constraint expected* scenarios than to unpredictable words in each of the four other conditions. Importantly, however, the N400 evoked by the four types of unpredictable words was not sensitive to the lexical constraint of the preceding context. Replicating Kutas & Hillyard (1984) and Federmeier et al. (2007) in a new stimulus set, the amplitude of the N400 evoked by critical words in the *high constraint unexpected* and the *low constraint unexpected* scenarios was indistinguishable. We further show that this insensitivity of the N400 to lexical constraint extended to anomalous critical words: the amplitude of the N400 to critical words in the *high constraint anomalous* and the *low constraint anomalous* scenarios was also indistinguishable. Taken together, these findings support the hypothesis that the N400 primarily reflects the degree of match between semantic features that are predicted by a context and the semantic features associated with an incoming word, rather than the detection or recovery from violations of strong lexico-semantic predictions.

In addition, we show that critical words whose semantic features mismatched animacy constraints of their preceding verbs (both in the *high constraint anomalous* and the *low constraint anomalous* scenarios) elicited a slightly larger N400 amplitude than unpredicted but plausible critical words that were consistent with these animacy constraints (*low constraint unexpected* and *high constraint unexpected* scenarios).^5^ This finding shows that the amplitude of the N400 can be sensitive to implausibility, over and above lexical predictability (as operationalized by cloze), and over and above the effects of simple semantic relationships between the critical word and the prior context (as operationalized by LSA). Rather than reflecting a post-lexical integrative mechanism, we interpret this N400 plausibility effect as arising from the same semantic predictive mechanism that gave rise to the N400 effects of cloze probability — a point that we return to below.

### Late positivities: Violations of strong predictions

In contrast to the N400, the late positive components were selectively modulated by *violations* of strong predictions. Moreover, the scalp topographies of these positivities differed depending on the grain of representation that was violated: Critical words that only violated specific event/lexical constraints without violating event structure/animacy constraints (*high constraint unexpected*) selectively evoked a *late frontal positivity*, which was larger than that produced by critical words in any other condition. In contrast, critical words that only violated event structure/animacy constraints (*low constraint anomalous*) evoked a *late posterior positivity/P600,* which was larger than that produced by critical words in any of the three plausible scenarios. These findings suggest that the underlying neural sources contributing to each of these effects are at least partially distinct.

Importantly, our findings on the double violations, which violated *both* event structure/animacy constraints and finer-grained event/lexical constraints, suggest that, although distinct, the neurocognitive mechanisms underlying the *late frontal positivity* and the *late posterior positivity/P600* are interdependent: The doubly violated *high constraint anomalous* critical words did not produce a *late frontal positivity* effect, and instead only produced a *late posterior positivity/P600* effect, which was larger than that produced by the *low constraint anomalous* critical words. This suggests a trade-off relationship between these two components, which has important implications for their functional interpretation.

With regard to the *late frontal positivity,* the absence of an effect on the doubly violated critical words suggests that this component cannot simply reflect the detection of a lexical prediction violation. In the *high constraint anomalous* scenarios, the critical word, “drawer”, also violated strong lexical constraints. If the *late frontal positivity* only reflected the detection of a discrepancy between the predicted and encountered lexical item, or a competitive process operating purely at the level of semantic features, then a frontal positivity should have also been evoked in this condition. The fact that no such effect was observed suggests that the *late frontal positivity* reflected a higher-level process that was engaged only in the *high constraint unexpected* scenarios. Specifically, we propose that this component indexes a large change in activity associated with successfully updating the higher-level situation model — the process of shifting from the prior situation model to a new situation model on the basis of new bottom-up input. As we discuss further below, we also suggest that this higher-level shift entailed top-down feedback suppression of the incorrectly predicted semantic features, and selection of the correct features.

With regard to the *late posterior positivity/P600*, which was selectively produced by the anomalous critical words, we similarly suggest that, rather than simply reflecting the detection of an animacy violation, this effect also reflected higher-level activity. Specifically, we suggest that it was triggered when the bottom-up input conflicted with the constraints of the existing *communication model*. This resulted in an (initial) *failure* to update the current situation model with the new input (see Shetreet et al., 2019 for recent discussion), thereby ‘blocking’ any *late frontal positivity* effect. Our finding that the *late posterior positivity/P600* was larger on the doubly violated critical words (*high constraint anomalous* > *low constraint anomalous*) suggests that the comprehender’s original high certainty prediction for a specific event made it easier to detect conflict (previous work suggests that the *detection* of anomalies is critical to producing the *late posterior positivity/P600*, see Sanford et al., 2011). This finding is consistent with previous reports that the *late posterior positivity/P600* evoked by syntactic violations is also influenced by the lexical constraint of the preceding context: its amplitude is larger to syntactic anomalies within high constraint than low constraint contexts (Gunter, Friederici, Schriefers, 2000).

As we discuss next, this pattern of ERP findings can be understood within a *hierarchical generative framework* of language comprehension (Kuperberg & Jaeger, 2016; Kuperberg, 2016; see also Xiang & Kuperberg, 2015). Below we outline this framework and offer functional interpretations of the N400, the *late frontal positivity* and the *late posterior positivity/P600* components within this framework. We then discuss these interpretations in relation to previous interpretations of these components, previous models of language comprehension, and more general accounts of predictive processing in the brain.

### A hierarchical generative framework of language comprehension

Within a hierarchical generative framework of language comprehension (see Kuperberg & Jaeger, 2016, section 5, page 16; Kuperberg, 2016), the agent draws upon an internal generative model^6^ — a hierarchical network of stored linguistic and non-linguistic representations that she believes are relevant to her communicative goals. At the top of this generative network is the *communication model,* which represents the comprehender’s beliefs about the communicator and the broader communicative environment. As noted in the Introduction, in the present study, we assume that the comprehender believes that the communicator is an English speaker who is communicating literally about events that are possible in the real world (see also Frank & Goodman, 2012; Degen, Tessler & Goodman, 2015). If the agent’s goal is deep language comprehension, then, within this communication model, she will construct a *situation model* of the current discourse. Because the linguistic input unfolds linearly over time, and because it takes time to pass new information up from lower to higher levels of representation, the comprehender cannot achieve the goal of comprehension all at once. However, if she has hypotheses at a higher level of representation, then she can test them by propagating probabilistic predictions down to lower levels of the hierarchy, thereby pre-activating information at these lower levels. Then, as new bottom-up information becomes available to each level of representation, any information that confirms these top-down predictions is ‘explained away’ (its processing is facilitated), while any information that remains unexplained is propagated up the generative hierarchy where it can be used to update hypotheses represented at higher levels.

In most situations, at any given point in time, the arrival of unexplained input at the high-level situation model will induce only a small change in its state; larger changes are accumulated gradually over time. Sometimes, however, a particular input may induce a large change in state at this level. For example, if at an earlier point in time, the comprehender had already settled, with high certainty, on a particular high-level message that she believed the communicator intended to convey, then the arrival of new unexplained information at the situation model will lead to a large change in state, corresponding to the large shift in the comprehender’s interpretation.

So, as long as the bottom-up input is compatible with the constraints of the communication model (and all the levels below it within the generative network), then, through incremental cycles of prediction and hypothesis updating, the comprehender should be able to ‘home in’ on the intended message with increasing certainty as the linguistic input progressively unfolds over time. However, if the comprehender receives unpredicted information that *conflicts* with the constraints of the communication model itself, then these cycles of updating the situation model come to a halt, leading to a temporary comprehension failure. This may trigger prolonged second-pass attempts to make sense of the discourse scenarios through reanalysis, attempts to repair the input, and/or attempts to revise the communication model itself. We now illustrate these principles with the example stimuli used in this study.

As shown in the top left panels of Figures 7 and 8, we assume that, after reading a *low constraint* context, but before encountering the critical word, the comprehender has established a simple situation model that constrains for the event structure, {*Agent cautioned animate-Patient*} (see also Altmann & Mirkovic, 2009). This leads to the prediction of semantic features that are characteristic of animate entities (e.g. <sentient>, <can move>). As shown in the bottom left panels of Figures 7 and 8, we assume that, after reading a *high constraint* context, the comprehender has built a rich situation model, leading her to strongly predict the specific event, {*Lifeguards cautioned Swimmers*}(see also Altmann & Mirkovic, 2009; Hare, Jones, Thomson, Kelly, & McRae, 2009; McRae & Matsuki, 2009). This, in turn, leads her to predict not only semantic features associated with animate entities (e.g. <sentient>, <can move>), but also additional semantic features associated with “swimmers” (e.g. <in water>, <afloat>).^7^

**Figure 7.**
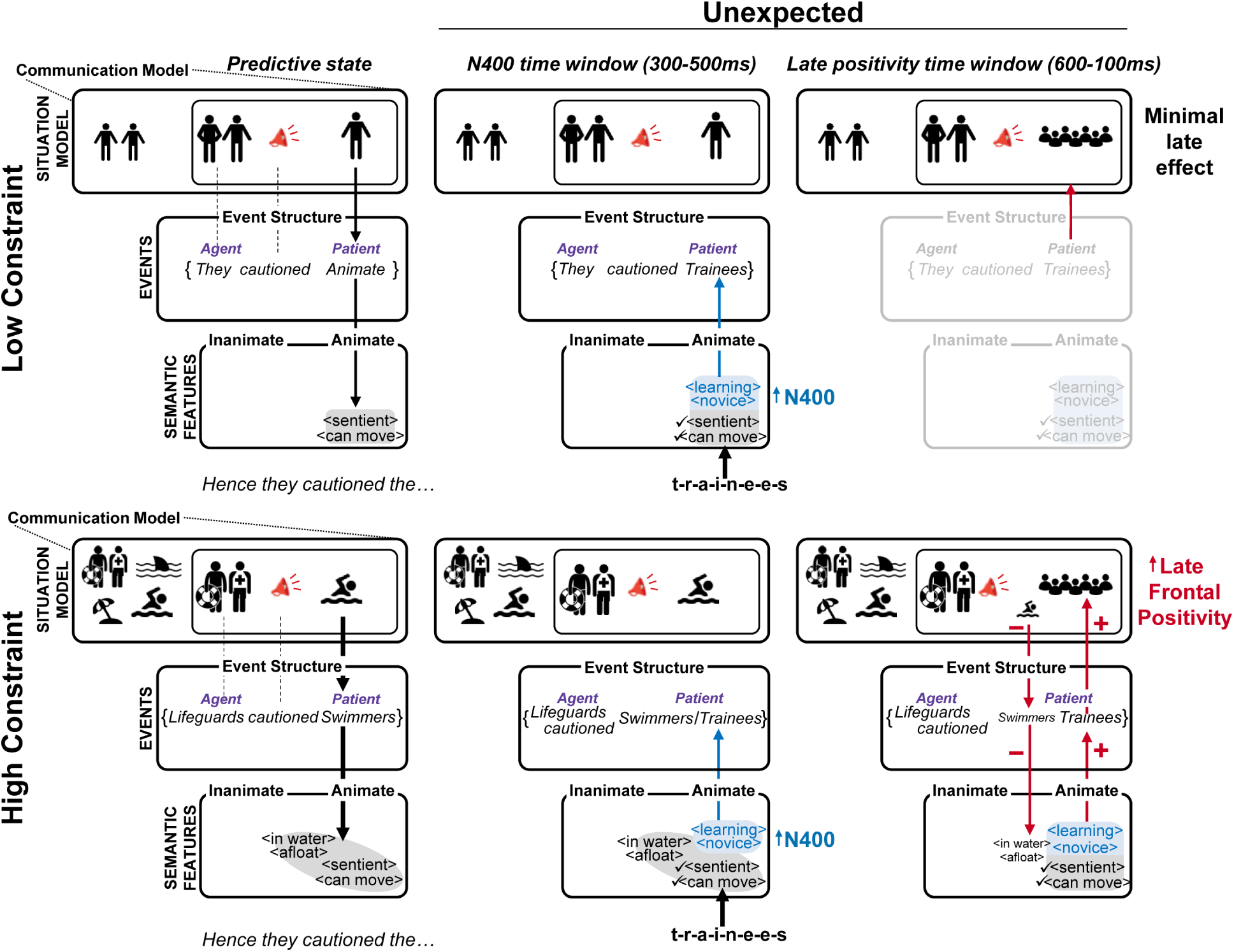
Schematic illustration of the state of the language comprehension system at three hierarchical levels of representation before and after encountering plausible critical nouns. States associated with the *low constraint unexpected* scenarios (Condition 2 in Table 1) are illustrated in the top half of the figure, and states associated with the *high constraint unexpected* scenarios (Condition 3 in Table 1) are illustrated in the bottom half of the figure. The left panels show the predictive before encountering the critical word; the middle panels show the state during the N400 time window (300-500ms); the right panels show the state during the late positivity time window (600-1000ms). Please see text pages 35-39 for full explanation.

**Figure 8.**
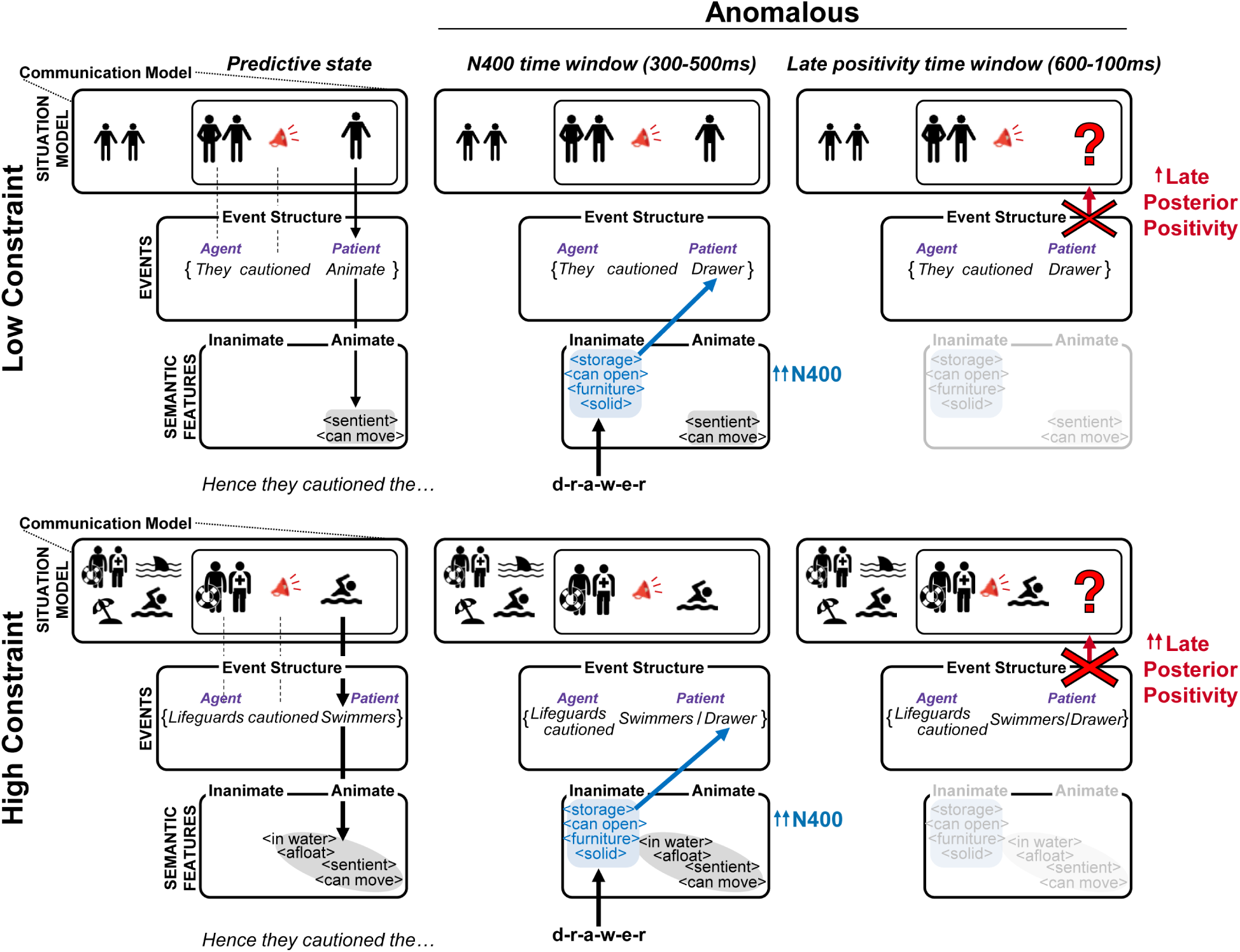
Schematic illustration of the state of the language comprehension system at three hierarchical levels of representation before and after encountering anomalous critical nouns. States associated with the *low constraint anomalous* scenarios (Condition 4 in Table 1) are illustrated in the top half of the figure, and states associated with the *high constraint anomalous* scenarios (Condition 5 in Table 1) are illustrated in the bottom half of the figure. The left panels show the predictive state before encountering the critical word; the middle panels show the state during the N400 time window (300-500ms); the right panels show the state during the late positivity time window (600-1000ms). Please see text pages 35, 37, and 39-40 for full explanation.

Within this framework, the amplitude of the N400 evoked by the incoming critical word can be naturally understood as reflecting the retrieval of, or access to, semantic features that have not already been predicted by the context (see Kuperberg, 2016, for further discussion). In the *high constraint expected* scenarios, the critical word, “swimmers”, offers no new semantic information and so it evokes a small amplitude N400. In the middle panel of Figure 7, in both the *low constraint unexpected* and the *high constraint unexpected* scenarios, the semantic features associated with the critical word, “trainees”, match these pre-activated semantic features that characterize its animacy (<sentient>, <can move>), but the comprehender must retrieve additional semantic features that more specifically characterize “trainees” (e.g. <learning>, <novice>). The amplitude of the N400, which reflects the retrieval of these unpredicted semantic features, is therefore larger (see Figure 7, middle panel).

Finally, in both the *low constraint anomalous* and the *high constraint anomalous* scenarios, no semantic features associated with the critical word, “drawer”, have been pre-activated, and so when the bottom-up input is encountered, the comprehender must retrieve all its properties, including its mismatching inanimate features (e.g. <storage>, <can open>, <furniture>, <solid>), see also Paczynski & Kuperberg, 2011, 2012. The amplitude of the N400 is therefore even larger.

As shown in both Figures 7 and 8 (middle panel), in all four types of unpredictable scenarios, any unpredicted semantic features are passed up to higher levels of representation where, at a slightly later stage of processing, activity differs depending on condition. In the *high constraint unexpected* scenarios, the arrival of unpredicted input at the situation model produces a *late frontal positivity* effect. We suggest that this effect primarily reflects a large change in activity associated with successfully updating the situation model from its prior state to a new state. We further suggest that this high-level shift is linked to the top-down suppression of competing lower-level incorrect lexico-semantic predictions, and the selection of the correct lexico-semantic features. Specifically, we propose that, prior to encountering the critical word, the comprehender had built a rich situation model and had used this model to strongly predict a specific event, {*Lifeguards cautioned Swimmers*}, as well as the semantic features associated with <swimmers>, see bottom left panel of Figure 7. Then, as the unpredicted semantic features associated with the critical word, “trainees” (<learning>, <novice>), arrive at the event level, activity begins to shift towards a new event, {*Lifeguards cautioned Trainees*}, see bottom middle panel of Figure 7. This unpredicted event information is, in turn, passed up so that the comprehender begins to update her situation model. However, in order to actually *complete* updating the situation model, the comprehender must suppress the original incorrectly predicted event, {Lifeguards cautioned Swimmers}, and the incorrectly predicted semantic features associated with “swimmers”, thereby ‘selecting’ the newly inferred event, {Lifeguards cautioned Trainees}, and the semantic features associated with “trainees”, see bottom right panel of Figure 7.

In the *low constraint unexpected* scenarios, no significant *late frontal positivity* effect is produced by critical words. We suggest that this is because any shift in activity induced by the unpredicted semantic features at the level of the situation model was much smaller. As shown in the top middle panel of Figure 7, as soon as the unpredicted semantic features associated with the critical word, “trainees”, reach the event level, the comprehender is able to shift activity to infer the event, {*They cautioned Trainees*}, within the N400 time window. This is because the event structure has already been correctly predicted and there is no competition at the level of semantic features. In other words, by fully accessing these unpredicted semantic features, the amount of additional work to infer the specific event is minimal. It is possible that there may be some small additional shift as the comprehender updates her situation model in the late positivity time window, but this change in activity is much smaller than in the *high constraint unexpected* scenarios.^8^

Finally, we suggest that the *late posterior positivity/P600* is triggered by an initial *failure* to update the situation model when new input conflicts with the constraints of the broader communication model. This may be followed by second-pass attempts to make sense of the discourse scenarios through reanalysis, attempts at repair of the input, and/or revision of the communication model. As shown in the left panels of Figure 8, in both the *high constraint* and *low constraint* contexts, the situation model constrains for a particular event structure, {*Agent cautioned animate-Patient*} — the set of events that is compatible with the broader communication model. As shown in the middle panels of Figure 8, when unpredicted semantic features associated with “drawer” arrive at the event level, this leads the comprehender to infer a different event structure, {*Agent cautioned inanimate-Patient*}. This unexplained information is passed up to the situation model. However, unlike in the *high constraint unexpected* scenarios, the input cannot be explained at this level (the conflict cannot initially be resolved), triggering the *late posterior positivity/P600*. Note that there is greater conflict in the *high constraint anomalous* scenarios than in the *low constraint anomalous* scenarios because the original predictions were stronger following the *high constraint* than the *low constraint* contexts.

### Relationship with other functional interpretations of these ERP components and other models of language processing

#### The N400

A classic debate about the N400 is whether it reflects ‘lexical access’ (e.g. Lau, Phillips, Poeppel, 2008) or combinatorial ‘integration’^9^ (e.g. Hagoort, Baggio, & Willems, 2009). Implicit in this debate is the assumption that ‘lexical access’ and ‘integration’ are independent of one another and serial in nature, such that the comprehender first needs to achieve full access to a unique lexical item before she can begin to integrate this item into its preceding context.

The generative framework outlined above makes no such assumption. According to this framework, the comprehender can predict an upcoming event structure or a specific event (and its associated semantic features) *before* a new word becomes available from the bottom-up input (see also Metusalem et al., 2012; Rabovsky, Hansen & McClelland, 2018). When the new word is encountered, so long as its semantic and syntactic features are consistent with the predicted event structure/event, little additional work is required to ‘integrate’ it. In other words, because semantic predictions are derived from higher-level event-level predictions, in a sense, some or all of a predicted word’s integration has already been completed, even prior to its occurrence (for further discussion of such “pre-updating”, see Kuperberg & Jaeger, 2016, section 4, pages 14-15). In cases when a word violates strong prior event/lexical predictions (as in the *high constraint unexpected* scenarios), the additional work required to integrate it into the current situation model should primarily be reflected in the amplitude of later, post-N400 components, as discussed further below.

One argument that some have raised in favor of an integration account of the N400 is that its amplitude can be sensitive to the implausibility of the resulting interpretation (although the N400 is not always sensitive to plausibility, see Kuperberg, 2016, and Shetreet et al., 2019, Supplementary Materials section 3 for a recent review and discussion). The assumption is that, having accessed a lexical item, it takes more combinatorial work to construct an implausible/incoherent proposition than to construct a plausible/coherent proposition (see Lau, Namyst, Fogel & Delgado, 2016; Nieuwland et al., in press). The amplitude of the N400 is, in part, taken to reflect this additional work. Because effects of plausibility/coherence on the N400 are sometimes observed over and above the effects of lexical probability (as indexed by cloze probability), it has also sometimes been assumed that any effects of implausibility do not reflect the consequences of prior prediction (e.g. Nieuwland et al., in press; van Berkum, Zwitserlood, Hagoort & Brown, 2003).

Once again, the generative framework described above makes no such assumptions. We suggest that any sensitivity of the N400 to implausibility, over and above the effects of cloze probability, can again reflect the retrieval of semantic features that have not already been predicted. This is because comprehenders can, in some cases, use their situation model to predict event structures and their associated semantic features, *even when* these predictions do not correspond to the pre-activation of specific individual lexical items. For example, we have argued that the slightly larger N400 produced by the unpredictable critical word, “drawer”, in the anomalous scenarios reflected the additional retrieval of inanimate features that were not pre-activated by the context (see Szewczyk & Schriefers, 2013 and Wang, Jensen, & Kuperberg, 2018, for evidence that semantic features linked to animacy can be pre-activated prior to an incoming word). Of course, it is possible that shifts in activity at higher levels of representation may begin within the N400 time window (e.g. Nieuwland et al., in press; Lau et al. 2016). Our primary point is that any effects of plausibility on the N400 itself can ultimately stem from the *same* semantic predictive mechanism that gives rise to the effects of cloze probability on the N400.

We also emphasize that the pre-activation of semantic features by a higher-level representation cannot simply be explained by passive ‘resonance’ or ‘priming’ between content words, as is assumed in some theories of discourse comprehension (e.g. Myers & O’Brien,1998), as well as some views of the N400 (e.g. Brouwer, Fitz & Hoeks, 2012). This is because, in the present study, semantic relationships between the critical words and the ‘bag of words’ in the preceding context were matched between the unexpected and the anomalous conditions using LSA (see also Kuperberg, Paczynski & Ditman, 2011, and see Kuperberg, 2016, for discussion). Rather, we attribute any pre-activation of semantic features related to the animacy of the incoming word to the prediction of the event structure, *{Agent cautioned animate-Patient}*. Finally, we emphasize that the pre-activation of semantic features by a higher-level representation does not necessarily or always correspond to the prediction of specific stored lexical entities (see also Federmeier & Kutas, 1999; Laszlo & Federmeier, 2009; Kutas & Federmeier, 2011).

#### Late frontal positivity

As noted in the Introduction, the *late frontal positivity* is classically produced by words that violate strong lexical constraints. As such, it has been linked to the inhibition of incorrectly predicted information (Federmeier et al., 2007; Kutas, 1993; Ness & Meltzer-Asscher, 2018) and/or the integration of the violating bottom-up input to reach a new higher-level interpretation (Federmeier, Kutas & Schul, 2010; DeLong, Quante & Kutas, 2014; Brothers, Swaab & Traxler, 2015). These processes could be assumed to be serial and therefore independent of one another. For example, one possible account is that, having computed a partial high-level representation of the context (e.g. {The little boy went out to fly his…}), and used this to predict a specific upcoming lexical item (“kite”), the comprehender must first inhibit “kite” before she can begin to integrate “plane” into her current contextual representation. The *late frontal positivity* effect would be assumed to reflect either the inhibition and/or the integrative process.

Within the hierarchical generative framework, however, these processes are fundamentally interrelated and they proceed in parallel. Specifically, as discussed above, after reading the high constraint context and before encountering the incoming word, the comprehender is assumed to have *already* updated her situation model such that it includes a representation of the lifeguards warning a group of swimmers (‘pre-updating’). Within this framework, the *late frontal positivity* corresponds to the large shift in activity associated with *updating* this situation model when new information is encountered (‘re-integration’), as well as the top-down feedback suppression of incorrectly predicted semantic features, which is necessary to complete this re-updating process.

#### Late posterior positivity/P600

Debates about the functional significance of the *late posterior positivity/P600* have come from many different perspectives.

In terms of its primary trigger, several authors have proposed that, in order to produce the effect, the comprehender must detect *conflict* between alternative representations that are computed during language comprehension (e.g. Kim & Osterhout, 2005; Kuperberg, 2007; van de Meerendonk, et al., 2009). Others have emphasized that, unlike the *late frontal positivity*, the *late posterior positivity/P600* is produced by words that yield highly implausible interpretations (DeLong et al., 2014; Kuperberg, 2013; Van Petten & Luka, 2012), and, in particular, by words that are, at least initially, perceived as *impossible/anomalous* during online comprehension (for discussion see Kuperberg, 2007, section 3.4; e.g. Kuperberg et al., 2003; van de Meerendonk et al., 2010; Paczynski & Kuperberg, 2012).

The interpretation that we have offered above attempts to bridge these accounts. We argue that the primary trigger of the *late posterior positivity/P600* is the detection of high-level conflict with the comprehender’s broader communication model. As a result, the comprehender cannot initially incorporate the new input into her current situation model, leading to the initial perception of incoherence/impossibility. This account makes two assumptions. The first is that, in order to produce a *late posterior positivity/P600,* the comprehender must have previously established a communication model that now conflicts with the new bottom-up input. For example, in the present study, we assume that the comprehender had already established a model of a literal English speaker who communicates about events that are possible in the real world (cf. Frank & Goodman, 2012; Degen, Tessler & Goodman, 2015). Therefore, the event, *{Lifeguards cautioned Drawer}* produced a *late posterior positivity/P600* because it conflicted with these constraints. If, however, the comprehender had previously established a communication model that allowed for the expression of fantasy world events, in which drawers and other inanimate objects could understand language, then the same event would have been interpreted as plausible, and no *late posterior positivity/P600* would have been produced (e.g. see Nieuwland & Van Berkum, 2006).

The second assumption is that, to trigger the *late posterior positivity/P600*, the initial failure to incorporate the input into the situation model occurs during fast, online comprehension. While the online detection of incoherence often patterns with offline judgments of impossibility (as in the current study), this may not always be the case. *Late posterior positivity/P600* effect are sometimes seen in sentences that are not judged to be impossible offline, but in which some temporary conflict with the existing communication model is detected during online processing. For example, a *late posterior positivity/P600* can be evoked by unexpected words in semantically reversible sentences, even though these sentences are generally judged offline to be implausible-but-not-impossible (e.g. “The restaurant owner forgot which waitress the customer had served…”: Chow, Smith, Lau & Phillips, 2015; Kolk, Chwilla, van Herten & Oor, 2003; see Kuperberg, 2016 for discussion). In addition, a *late posterior positivity/P600* is sometimes evoked in non-literal constructions such as nominal metaphors (e.g. De Grauwe, Swain, Holcomb, Ditman & Kuperberg, 2010, Experiment 2) or in association with certain types of metonymic shifts (e.g. Schumacher, 2013).

In terms of the neurocognitive mechanisms reflected by the *late posterior positivity/P600,* one possibility is it *only* reflects the detection of high-level conflict, and the resulting initial failure to incorporate the input into the current situation model. Another possibility is that it additionally reflects subsequent prolonged attempts to make sense of the input. These might include a reanalysis of the prior context in order to check whether or not it was accurately perceived the first time around (van de Meerendonk et al., 2010), and, if necessary, attempts to repair it. Alternatively, it may reflect second-pass attempts to come up with a revised interpretation of the input (Kuperberg et al., 2006; Kuperberg, 2007; see also Brouwer, Fitz & Hoeks, 2012)). Within the generative framework described above, this type of re-interpretation would entail revising the constraints of the communication model itself (modifying its parameters) so that it can accommodate the previously impossible interpretation. For example, revising the communication model may allow the comprehender to accept a fantasy world scenario as possible, or to come to a non-literal interpretation of the input (e.g. Schumacher, 2013; De Grauwe, Swain, Holcomb, Ditman & Kuperberg, 2010). As we discuss further below, this type of revision of the communication model is closely linked to longer-term learning/adaptation.

From the current study alone, we cannot determine whether the *late posterior positivity/P600* effect reflects any of these additional processes. While our comprehension and judgment tasks suggest that participants were reading carefully for comprehension and successfully detecting the presence of semantic anomalies, they do not provide direct evidence that they were additionally engaged in processes related to re-analysis, repair, or re-interpretation. It will therefore be important for future experiments to further investigate the neurocognitive function of the *late posterior positivity/P600* by using tasks that explicitly ask participants to repair the input or attempt to create representations of literally impossible events.

Finally, we note that the *late posterior positivity/P600* is likely to share basic computational information processing mechanisms with the well-known posterior P300 component (Coulson, King & Kutas, 1998; Osterhout, Kim & Kuperberg, 2012; Sassenhagen, Schlesewsky & Bornkessel-Schlesewsky, 2014; Sassenhagen & Fiebach, 2019). We discuss these relationships in detail in Kuperberg & Brothers (in preparation).

### Relationship with more general theories of prediction, prediction error and predictive coding in the brain

The generative framework of language comprehension outlined above is keeping with more general theories that emphasize a central role of prediction in the brain (e.g. Clark, 2013), as well as with many previous studies showing that the brain can encode ‘prediction error’ — the difference between a predicted state of activity and the state after new input is observed (den Ouden, Kok & de Lange, 2012; Schutz & Dickinson, 2000).

Both probabilistic prediction and prediction error are important components of ‘predictive coding’ (Mumford, 1992; Rao & Ballard, 1999; Rao & Ballard, 1997; Friston, 2005), which has been proposed as an algorithm for carrying out Bayesian inference in the brain (although it is important to note that not all neural prediction error is in the service of Bayesian inference, and predictive coding may not be the only way in which Bayesian inference is carried out in the brain, see Lengyel & Aitchison, 2017). Hierarchical predictive coding involves passing up prediction error from lower to higher levels of the cortical hierarchy, updating beliefs at higher levels, and passing down new predictions to lower levels, with the overarching goal of minimizing prediction error across the entire cortical network (Friston, 2005). While we think that it is premature to conclude that language comprehension in the brain is carried out through predictive coding, we do note that there are features of our model that are consistent with some of its principles.

First, we have suggested that unpredicted semantic features, reflected by the N400, are passed up to higher levels of representation. This is consistent with the basic principle that prediction error at lower levels is passed up a generative hierarchy, and that it can be indexed by ERPs produced by superficial pyramidal cells (Friston, 2005). Importantly, here we assume that the ‘prediction error’, indexed by the N400, is at the level of semantic features, and that it can be formalized simply as a word’s semantic probability, independent of the probability of any other words that may have also been predicted by the prior context. In other words, when used in relation to the N400, the term, *prediction error* does *not* necessarily correspond to how the term, ‘prediction violation’ (or “error”) is typically used in psycholinguistic models.

Second, we have argued that the *late frontal positivity* reflects a high-level shift at the level of the situation model, which is induced by the arrival of unpredicted information that cannot be explained at lower levels of the generative hierarchy. This is consistent with the idea that, by passing up lower-level prediction error to higher levels of representation, higher-level prediction error can be reduced through Bayesian inference. The high-level shift of the situation model may correspond to *Bayesian surprise* (Baldi & Itti, 2010) --- the difference between the prior and the posterior probability distribution of higher-level hypotheses. Moreover, we have also suggested that this high-level shift is accompanied by top-down feedback suppression/selection to lower levels of representation. This is consistent with some versions of predictive coding which propose that, having corrected higher-level hypotheses, earlier incorrect lower level predictions can also be retrospectively corrected through feedback suppression. This, in turn, leads to a relative enhancement of stimulus-driven activity that is consistent with the new higher-level hypotheses (see Lee & Mumford, 2003; Spratling, 2008).

Finally, we have argued that the *late posterior positivity/P600* is triggered by a failure to incorporate unpredicted input into the existing situation model because of conflict with the communication model. This can be conceptualized as reflecting unresolved prediction error at the highest level of the generative hierarchy. The prediction error is unresolved because there is a discrepancy between the input and the model that the comprehender had previously been assumed (cf. ‘puzzlement’ surprise, see Faraji, Preuschoff & Gerstner 2018). As discussed briefly below, this ‘unexpected surprise’ (Yu & Dayan, 2005) might cue the comprehender to revise the communication model, or to switch models (adaptation), again with the broad goal of reducing overall prediction error.

### Implications

By associating different ERP components with neural activity at different levels of a hierarchy of representations, this hierarchical generative framework makes several predictions and has several implications.

First, to the degree that different hierarchical levels of representation map on to neuroanatomically distinct cortical regions, this framework predicts not only a temporal segregation of responses to words that confirm and violate predictions, but also some spatial segregation of neural activity. Consistent with this hypothesis, we have recently carried out a multimodal neuroimaging study, using ERP together with MEG and fMRI, to show that, within the N400 time window, confirmed predictions are associated with reduced activity within the left temporal cortex, while prediction violations are associated with additional recruitment of the left inferior prefrontal cortex, with feedback activity to different parts of the temporal cortex depending on the grain of representation that was violated.

Second, the generative hierarchical framework predicts that, under certain circumstances, it should be possible to violate lower level predictions without seeing evidence of a higher-level shift at the level of the situation model. For example, if comprehenders are engaged in shallow processing, and have not established a higher-level situation model, then lexical prediction violations may be associated with a large N400, without triggering a *late frontal positivity*, and animacy violations may be associated with a large N400 without triggering a *late posterior positivity/P600* (see Brothers, Wlotko, Warnke & Kuperberg, under review).

Conversely, it should also be possible to see evidence of higher-level activity without violating lower-level predictions. Some evidence for this claim comes from studies reporting a *late frontal positivity* in response to words that do not violate strong lexical constraints of their preceding contexts (e.g. Chow, Lau, Wang & Phillips, 2018; Freunberger & Roehm, 2016; Thornhill & Van Petten, 2012; Zirnstein, van Hell & Kroll, 2018;). These “low-constraint” frontal positivities may occur if a new input to the situation model is particularly informative, triggering a large update, even when the prior situation model had not led to strong lower-level predictions of upcoming semantic features. Similarly a *late posterior positivity/P600* can sometimes be evoked by words that do not violate strong lower-level animacy or other selection restriction if the bottom-up input conflicts with presuppositions that have previously been encoded and maintained within the comprehender’s high-level situation model (e.g. Shetreet et al., 2019).

We also emphasize that the precise conditions evoking the late positivities are likely to vary across languages. For example, in some languages (e.g. Spanish, Dutch), a mismatching gender-marked adjective or determiner may provide strong evidence that a predicted specific event will be violated, leading the comprehender to update her situation model and produce a *late frontal positivity*, even before she receives direct evidence of violating semantic features (e.g. Wicha, Moreno & Kutas, 2004). Moreover, conditions evoking the late positivities are also likely to vary between individuals, where variability in linguistic and domain-general cognitive abilities may predict differences in the amplitude of these components (e.g. see Zirnstein, van Hell & Kroll, 2018 for evidence of individual variability in the *late frontal positivity;* see Nakano, Saron & Swaab, 2010 and Kim, Oines & Miyake, 2018 for evidence of individual variability in the *late posterior positivity/P600*).

Finally, within this generative framework, we can begin to understand *why* the brain should engage different neurocognitive mechanisms in response to predictions that are confirmed versus predictions that are strongly violated. This is because the framework explicitly links the process of language *comprehension* to language *adaptation* (see also Kleinschmidt & Jaeger, 2015; Chang, Dell, & Bock, 2006; Dell & Chang, 2014). In the present study, we have focused on comprehension, discussing the late positivities as responses that can potentially help comprehenders to recover meaning when input violates strong predictions, by updating the current situation model (the *late frontal positivity*) or by diagnosing that something is wrong with the input and reanalyzing/repairing/reinterpreting it (the *late posterior positivity/P600*). It is sometimes assumed that these types of neurocognitive mechanisms reflect unwanted *costs* of a predictive language comprehension system (e.g. Federmeier, 2007, and Kutas, DeLong, & Smith, 2011 for discussion). However, another way of understanding their *broader* functional role is as providing strong signals that the statistical structure of the broader communicative environment has *changed ––* so-called ‘unexpected surprise’ (Yu & Dayan, 2005). This type of signal may cue the comprehender to *adapt* to her new communicative environment, which is necessary for her to continue predicting efficiently. Within the hierarchical generative framework discussed here, adaptation would involve modifying the *parameters* of the comprehender’s existing generative network. This may entail revising the existing communication model so that it can accommodate future fantasy world scenarios, or metaphorical interpretations (as discussed above). It may also entail revising assumptions about the reliability of the communicator, leading to a systematic down-weighting of linguistic cues at lower levels of the generative network. Alternatively, it might involve switching to a new generative network (cf Qian, Jaeger & Aslin, 2012, 2016; Gallistel, Krishan, Liu, Miller & Latham, 2014; see Kleinschmidt & Jaeger, 2015 for a detailed discussion). Some existing evidence suggests that the *late posterior positivity/P600* evoked by syntactic anomalies adapts over time (Coulson, King & Kutas, 1998; Hanulíková, Alphen, van Goch & Weber, 2012; see also Kuperberg & Brothers, in preparation, for discussion). It will be important for future studies to determine whether the late positivities play a functional role in actually driving such adaptation processes.

## Acknowledgments

This work was funded by the National Institute of Child Health and Human Development (R01 HD08252 to G.R.K.). E.W.W. was supported by an Institutional Research and Academic Career Development Award from the National Institute of General Medical Sciences (K12GM074869 to Tufts University, PI C. Moore). We thank Maria Luiza Cunha Lima, Margarita Zeitlin, and Connie Choi for their contributions to constructing the experimental materials, Margarita Zeitlin and Simone Riley for their assistance with data collection, Sophie Greene for her help in statistical analysis, and Lotte Schoot and Lin Wang for their insightful comments on the manuscript. We are very grateful to Arim Choi Perrachione for her tremendous patience and help in making (and re-making) the figures.

## Footnotes

1 ERP effects of contextual predictability can sometimes appear to begin before 300ms. It has been hypothesized that this earlier divergence reflects an ERP component that is distinct from the N400 and that is more sensitive to the effects of predicting phonological/orthographic properties of an incoming word (e.g. Lau, Holcomb & Kuperberg, 2013; Brothers, Swaab, & Traxler, 2015).

2 In the present study, we focus on the semantic P600. While some of the ideas we propose are relevant to understanding the syntactic P600 (as well as posteriorly distributed late positivities that are evoked by other types of linguistic and non-linguistic violations), a full discussion of these relationships is outside the scope of this paper.

3 In the present study, the event structure was defined largely by the thematic properties of the verb in the third sentence — properties that define the *semantic roles* around specific types of actions or states (Dowty, 1989; Gruber, 1965; Fillmore, 1967; Jackendoff, 1987). However, other types of linguistic cues can also constrain strongly for particular sets of events (event structures), e.g. presupposition triggers (see Shetreet, Alexander, Romoli, Chierchia & Kuperberg, 2019) and concessive discourse connectives (see Xiang & Kuperberg, 2015).

4 We initially carried out statistical analyses on trial-averaged data within each of these spatiotemporal regions of interest. However, as a reviewer pointed out, this limits our ability to generalize our findings to new sets of items (see Clark, 1973). The pattern of results for these two sets of analyses did not differ. We also carried out an initial omnibus ANOVA analyses on the trial-averaged data, with Scenario Type (five levels corresponding to the five conditions) and Region (five levels corresponding to five regions across the anterior-posterior distribution of the scalp) as within-subjects variables. These analyses confirmed that both the N400 and the late positivity effects differed significantly across the five conditions in their scalp distribution (significant Scenario Type by Region interactions in both the 300-500ms and the 600-1000ms time windows, *F*s > 9.66, *p*s < .0001).

5 Similar to previous findings (Paczynski & Kuperberg, 2011), the degree of N400 modulation depended on whether the verb constrained for an animate or inanimate direct object, and on the animacy of the argument. A detailed discussion of these interactions is beyond the scope of the current paper.

6 Generative models are classically described within Bayesian probabilistic frameworks of cognition (Griffiths, Kemp & Tenenbaum, 2008) at Marr’s first computational level of analysis. However, they can also be instantiated at Marr’s second algorithmic level (see McClelland 1998 & 2013).

7 A generative framework also assumes that, at least under some circumstances, the comprehender can predict/pre-activate information at other levels of representation. In the present study, this would include other lexical properties associated with the upcoming critical word. Specifically, in both the *high constraint* and *low constraint* contexts, she is likely to have predicted its syntactic representation (a noun-phrase). And, in the *high constraint* contexts, she may have also predicted its phonological/orthographic representation (*swimmers*).

8 Although the *late frontal positivity* effect evoked by the *low constraint unexpected* (versus the *high constraint expected*) critical words was not significant, examination of the voltage map for this contrast (shown in Figure 6, left panel) suggests that the spatial distribution of this effect was qualitatively similar to the distribution of the effect observed when contrasting the *high constraint unexpected* and the *high constraint expected* critical words within this time window.

9 Note that the term ‘integration’ in relation to the N400 has *not* always been used to imply combinatorial processing. It has sometimes been used simply to refer to the use of context to facilitate semantic processing of incoming words, only once some bottom-up information becomes available, without any assumption that this entails combinatorial processes that build new propositional meaning (e.g. Chwilla, Hagoort & Brown, 1998; Van Petten & Luka, 2012).

## References

Altmann, G. T., & Mirkovic, J. (2009). Incrementality and prediction in human sentence processing. Cogn Sci, 33(4), 583–609. doi:10.1111/j.1551-6709.2009.01022.x

Baldi, P., & Itti, L. (2010). Of bits and wows: A Bayesian theory of surprise with applications to attention. Neural Netw, 23(5), 649–666. doi:10.1016/j.neunet.2009.12.007

Balota, D. A., Yap, M. J., Hutchison, K. A., Cortese, M. J., Kessler, B., Loftis, B., . . . Treiman, R. (2007). The English lexicon project. Behav Res Methods, 39(3), 445–459.

Barr, D. J., Levy, R., Scheepers, C., & Tily, H. J. (2013). Random effects structure for confirmatory hypothesis testing: Keep it maximal. Journal of Memory and Language, 68(3), 255–278. doi:10.1016/j.jml.2012.11.001

Bates, D. M., Mächler, M., Bolker, B., & Walker, S. (2015). Fitting linear mixed-effects models using lme4. Journal of Statistical Software, 67(1), 1–48. doi:10.18637/jss.v067.i01

Brothers, T., Swaab, T. Y., & Traxler, M. J. (2015). Effects of prediction and contextual support on lexical processing: prediction takes precedence. Cognition, 136, 135–149. doi:10.1016/j.cognition.2014.10.017

Brothers, T., Wlotko, E. W., Warnke, L., & Kuperberg, G. R. (under review). Going the extra mile: Effects of discourse context on two late positivities during language comprehension.

Brouwer, H., Fitz, H., & Hoeks, J. (2012). Getting real about semantic illusions: Rethinking the functional role of the P600 in language comprehension. Brain Research, 1446, 127–143. doi:10.1016/J.Brainres.2012.01.055

Chang, F., Dell, G. S., & Bock, J. K. (2006). Becoming syntactic. Psychological Review, 113(2), 234–272. doi:10.1037/0033-295x.113.2.234

Chow, W. Y., Lau, E. F., Wang, S., & Phillips, C. (2018). Wait a second! delayed impact of argument roles on on-line verb prediction. Language, Cognition and Neuroscience, 33(7), 803–828. doi:10.1080/23273798.2018.1427878

Chow, W. Y., Smith, C., Lau, E. F., & Phillips, C. (2015). A “bag-of-arguments” mechanism for initial verb predictions. Language, Cognition and Neuroscience, 31(5), 577–596.

Chwilla, D. J., Hagoort, P., & Brown, C. M. (1998). The mechanism underlying backward priming in a lexical decision task: Spreading activation versus semantic matching. Quarterly Journal of Experimental Psychology Section A: Human Experimental Psychology, 51(3), 531–560.

Clark, A. (2013). Whatever next? Predictive brains, situated agents, and the future of cognitive science. Behavioral and Brain Sciences, 36(3), 181–204. doi:10.1017/S0140525X12000477

Clark, H. H. (1973). The language-as-a-fixed-effect fallacy: A critique of language statistics in psychological research. Journal of Verbal Learning and Verbal Behavior, 12(4), 335–359. doi:10.1016/S0022-5371(73)80014-3

Coulson, S., King, J. W., & Kutas, M. (1998). Expect the unexpected: Event-related brain responses to morphosyntactic violations. Language and Cognitive Processes, 13(1), 21–58. doi:10.1080/016909698386582

De Grauwe, S., Swain, A., Holcomb, P. J., Ditman, T., & Kuperberg, G. R. (2010). Electrophysiological insights into the processing of nominal metaphors. Neuropsychologia, 48(7), 1965–1984.

Degen, J., Tessler, M. H., & Goodman, N. D. (2015). Wonky worlds: Listeners revise world knowledge when utterances are odd. Presented at the Proceedings of the Thirty-Seventh Annual Conference of the Cognitive Science Society.

Dell, G. S., & Chang, F. (2014). The P-chain: relating sentence production and its disorders to comprehension and acquisition. Philosophical Transactions of the Royal Society B: Biological Sciences, 369(1634), 20120394. doi:10.1098/rstb.2012.0394

DeLong, K. A., Quante, L., & Kutas, M. (2014). Predictability, plausibility, and two late ERP positivities during written sentence comprehension. Neuropsychologia, 61C, 150–162. doi:10.1016/j.neuropsychologia.2014.06.016

DeLong, K. A., Urbach, T. P., Groppe, D. M., & Kutas, M. (2011). Overlapping dual ERP responses to low cloze probability sentence continuations. Psychophysiology, 48(9), 1203–1207. doi:10.1111/j.1469-8986.2011.01199.x

Delorme, A., & Makeig, S. (2004). EEGLAB: an open source toolbox for analysis of single-trial EEG dynamics including independent component analysis. J Neurosci Methods, 134(1), 9–21. doi:10.1016/J.Jneumeth.2003.10.009

den Ouden, H. E., Kok, P., & de Lange, F. P. (2012). How prediction errors shape perception, attention, and motivation. Frontiers In Psychology, 3, 548. doi:10.3389/fpsyg.2012.00548

Dowty, D. R. (1989). On the semantic content of the notion of thematic role. In G. Cherchia, B. Partee, & R. Turner (Eds.), Properties, Types and Meaning (pp. 69–129). Norwell, MA: Kluwer.

Faraji, M., Preuschoff, K., & Gerstner, W. (2018). Balancing New against Old Information: The Role of Puzzlement Surprise in Learning. Neural Computation, 30(1), 34–83. doi:10.1162/neco_a_01025

Federmeier, K. D. (2007). Thinking ahead: the role and roots of prediction in language comprehension. Psychophysiology, 44(4), 491–505. doi:10.1111/j.1469-8986.2007.00531.x

Federmeier, K. D., & Kutas, M. (1999). A rose by any other name: Long-term memory structure and sentence processing. Journal of Memory and Language, 41(4), 469–495. doi:10.1006/Jmla.1999.2660

Federmeier, K. D., Kutas, M., & Schul, R. (2010). Age-related and individual differences in the use of prediction during language comprehension. Brain Lang, 115(3), 149–161. doi:10.1016/j.bandl.2010.07.006

Federmeier, K. D., Wlotko, E. W., De Ochoa-Dewald, E., & Kutas, M. (2007). Multiple effects of sentential constraint on word processing. Brain Research, 1146, 75–84. doi:10.1016/j.brainres.2006.06.101

Fillmore, C. J. (1967, 1967). The case for case. Presented at the Texas Symposium on Language Universals.

Frank, M. C., & Goodman, N. D. (2012). Predicting pragmatic reasoning in language games. Science, 336(6084), 998. doi:10.1126/science.1218633

Freunberger, D., & Roehm, D. (2016). Semantic prediction in language comprehension: evidence from brain potentials. Language, Cognition and Neuroscience, 31(9), 1193–1205.

Friston, K. J. (2005). A theory of cortical responses. Philosophical Transactions of the Royal Society B: Biological Sciences, 360(1456), 815–836. doi:10.1098/Rstb.2005.1622

Gallistel, C. R., Krishan, M., Liu, Y., Miller, R., & Latham, P. E. (2014). The perception of probability. Psychological Review, 121(1), 96.

Griffiths, T. L., Kemp, C., & Tenenbaum, J. B. (2008). Bayesian models of cognition. In R. Sun (Ed.), The Cambridge Handbook of Computational Psychology (pp. 59–100). New York: Cambridge University Press.

Gruber, J. S. (1965). Studies in lexical relations. (Doctoral dissertation), Massachusetts Institute of Technology,

Gunter, T. C., Friederici, A. D., & Schriefers, H. (2000). Syntactic gender and semantic expectancy: ERPs reveal early autonomy and late interaction. J Cogn Neurosci, 12(4), 556–568.

Hagoort, P., Baggio, G., & Willems, R. M. (2009). Semantic unification. In M. S. Gazzaniga (Ed.), The Cognitive Neurosciences, 4th Ed. (pp. 819–836). Cambridge, MA: MIT Press.

Hagoort, P., Brown, C., & Groothusen, J. (1993). The syntactic positive shift (SPS) as an ERP measure of syntactic processing. Language and Cognitive Processes, 8(4), 439–483. doi:10.1080/01690969308407585

Hanulikova, A., van Alphen, P. M., van Goch, M. M., & Weber, A. (2012). When one person’s mistake is another’s standard usage: The effect of foreign accent on syntactic processing. J Cogn Neurosci, 24(4), 878–887. doi:10.1162/jocn_a_00103

Hare, M., Jones, M., Thomson, C., Kelly, S., & McRae, K. (2009). Activating event knowledge. Cognition, 111(2), 151–167. doi:10.1016/j.cognition.2009.01.009

Jackendoff, R. (1987). The status of thematic relations in linguistic theory. Linguistic Inquiry, 18(3), 369–411.

Kim, A., Oines, L., & Miyake, A. (2018). Individual differences in verbal working memory underlie a tradeoff between semantic and structural processing difficulty during language comprehension: An ERP investigation. *Journal of Experimental Psychology: Learning*, Memory and Cognition, 44(3), 406–420. doi:10.1037/xlm0000457

Kim, A., & Osterhout, L. (2005). The independence of combinatory semantic processing: Evidence from event-related potentials. Journal of Memory and Language, 52(2), 205–225. doi:10.1016/J.Jml.2004.10.002

Kleinschmidt, D. F., & Jaeger, F. T. (2015). Robust speech perception: Recognize the familiar, generalize to the similar, and adapt to the novel. Psychol Rev, 122(2), 148–203. doi:10.1037/a0038695

Kolk, H. H. J., Chwilla, D. J., van Herten, M., & Oor, P. J. (2003). Structure and limited capacity in verbal working memory: A study with event-related potentials. Brain Lang, 85(1), 1–36. doi:10.1016/S0093-934X(02)00548-5

Kuperberg, G. R. (2007). Neural mechanisms of language comprehension: Challenges to syntax. Brain Research, 1146, 23–49. doi:10.1016/j.brainres.2006.12.063

Kuperberg, G. R. (2013). The proactive comprehender: What event-related potentials tell us about the dynamics of reading comprehension. In B. Miller, L. Cutting, & P. McCardle (Eds.), Unraveling Reading Comprehension: Behavioral, Neurobiological, and Genetic Components (pp. 176–192). Baltimore, MD: Paul Brookes Publishing.

Kuperberg, G. R. (2016). Separate streams or probabilistic inference? What the N400 can tell us about the comprehension of events. Language, Cognition and Neuroscience, 31(5), 602–616.

Kuperberg, G. R., & Brothers, T. (in preparation). What can the P300 tell us about the P600? Understanding language comprehension within a decision theoretic framework.

Kuperberg, G. R., Caplan, D., Sitnikova, T., Eddy, M., & Holcomb, P. J. (2006). Neural correlates of processing syntactic, semantic, and thematic relationships in sentences. Language and Cognitive Processes, 21(5), 489–530. doi:10.1080/01690960500094279

Kuperberg, G. R., & Jaeger, T. F. (2016). What do we mean by prediction in language comprehension? Language, Cognition and Neuroscience, 31(1), 32–59. doi:10.1080/23273798.2015.1102299

Kuperberg, G. R., Kreher, D. A., Sitnikova, T., Caplan, D. N., & Holcomb, P. J. (2007). The role of animacy and thematic relationships in processing active English sentences: evidence from event-related potentials. Brain Lang, 100(3), 223–237. doi:10.1016/j.bandl.2005.12.006

Kuperberg, G. R., Paczynski, M., & Ditman, T. (2011). Establishing causal coherence across sentences: an ERP study. Journal of Cognitive Neuroscience, 23(5), 1230–1246. doi:10.1162/jocn.2010.21452

Kuperberg, G. R., Sitnikova, T., Caplan, D., & Holcomb, P. J. (2003). Electrophysiological distinctions in processing conceptual relationships within simple sentences. Cognitive Brain Research, 17(1), 117–129. doi:10.1016/S0926-6410(03)00086-7

Kutas, M. (1993). In the company of other words: Electrophysiological evidence for single-word and sentence context effects. Language and Cognitive Processes, 8, 533–572.

Kutas, M., DeLong, K. A., & Smith, N. J. (2011). A look around at what lies ahead: Prediction and predictability in language processing. In M. Bar (Ed.), Predictions in the brain: Using our past to generate a future (pp. 190–207). New York: Oxford University Press.

Kutas, M., & Federmeier, K. D. (2011). Thirty years and counting: finding meaning in the N400 component of the event-related brain potential (ERP). Annual Review of Psychology, 62, 621–647. doi:10.1146/annurev.psych.093008.131123

Kutas, M., & Hillyard, S. A. (1984). Brain potentials during reading reflect word expectancy and semantic association. Nature, 307(5947), 161–163. doi:10.1038/307161a0

Kuznetsova, A., Brockhoff, P. B., & Christensen, R. H. B. (2015). Tests for random and fixed effects for linear mixed effect models (lmer objects of lme4 package). R package version 2.0-33. http://cran.r-project.org/package=lmerTest.

Landauer, T. K., & Dumais, S. T. (1997). A solution to Plato’s problem: The Latent Semantic Analysis theory of acquisition, induction, and representation of knowledge. Psychol Rev, 104(2), 211–240. doi:10.1037/0033-295x.104.2.211

Landauer, T. K., Foltz, P. W., & Laham, D. (1998). An introduction to Latent Semantic Analysis. Discourse Process, 25(2-3), 259–284. doi:10.1080/01638539809545028

Laszlo, S., & Federmeier, K. D. (2009). A beautiful day in the neighborhood: An event-related potential study of lexical relationships and prediction in context. Journal of Memory and Language, 61(3), 326–338. doi:10.1016/j.jml.2009.06.004

Lau, E. F., Holcomb, P. J., & Kuperberg, G. R. (2013). Dissociating N400 effects of prediction from association in single-word contexts. J Cogn Neurosci, 25(3), 484–502. doi:10.1162/jocn_a_00328

Lau, E. F., Namyst, A., Fogel, A., & Delgado, T. (2016). A direct comparison of N400 effects of predictability and incongruity in adjective-noun combination. Collabra, 2(1). doi:10.1525/collabra.40

Lau, E. F., Phillips, C., & Poeppel, D. (2008). A cortical network for semantics: (De)constructing the N400. Nature Reviews Neuroscience, 9(12), 920–933. doi:10.1038/nrn2532

Lee, T. S., & Mumford, D. (2003). Hierarchical Bayesian inference in the visual cortex. Journal of the Optical Society of America A, 20(7), 1434. doi:10.1364/josaa.20.001434

Lengyel, M., & Aitchison, L. (2017). With or without you: predictive coding and Bayesian inference in the brain. Current Opinion in Neurobiology, 46, 219–227.

Levin, B. (1993). English Verb Classes And Alternations: A Preliminary Investigation. Chicago: University of Chicago Press.

Lopez-Calderon, J., & Luck, S. J. (2014). ERPLAB: an open-source toolbox for the analysis of event-related potentials. Frontiers in Human Neuroscience, 8, 213. doi:10.3389/fnhum.2014.00213

Lund, K., & Burgess, C. (1996). Producing high-dimensional semantic spaces from lexical co-occurrence. Behavior Research Methods, Instruments, & Computers, 28(2), 203–208.

McClelland, J. L. (1998). Connectionist models and Bayesian inference. In M. Oaksford & N. Chater (Eds.), Rational Models of Cognition (pp. 21–52). New York: Oxford University Press.

McClelland, J. L. (2013). Integrating probabilistic models of perception and interactive neural networks: a historical and tutorial review. Front Psychol, 4, 503. doi:10.3389/fpsyg.2013.00503

McRae, K., & Matsuki, K. (2009). People use their knowledge of common events to understand language, and do so as quickly as possible. Language and Linguistics Compass, 3(6), 1417–1429. doi:10.1111/j.1749-818X.2009.00174.x

Metusalem, R., Kutas, M., Urbach, T. P., Hare, M., McRae, K., & Elman, J. L. (2012). Generalized event knowledge activation during online sentence comprehension. Journal of Memory and Language, 66(4), 545–567. doi:10.1016/j.jml.2012.01.001

Morey, R. D. (2008). Confidence intervals from normalized data: A correction to Cousineau (2005). Tutorial in Quantitative Methods for Psychology, 4(2), 61–64.

Mumford, D. (1992). On the computational architecture of the neocortex. II. The role of cortico-cortical loops. Biological Cybernetics, 66(3), 241–251.

Myers, J. L., & O’Brien, E. J. (1998). Accessing the discourse representation during reading. Discourse Processes, 26(2&3), 131–157. doi:10.1080/01638539809545042

Nakano, H., Saron, C., & Swaab, T. Y. (2010). Speech and span: working memory capacity impacts the use of animacy but not of world knowledge during spoken sentence comprehension. Journal of Cognitive Neuroscience, 22(12), 2886–2898. doi:10.1162/jocn.2009.21400

Ness, T., & Meltzer-Asscher, A. (2018). Lexical inhibition due to failed prediction: Behavioral evidence and ERP correlates. *Journal of Experimental Psychology: Learning*, Memory and Cognition, 44(8), 1269–1285. doi:10.1037/xlm0000525

Nieuwland, M. S., Barr, D. J., Bartolozzi, F., Busch-Moreno, S., Darley, E., Donaldson, D. I., . . . Von Grebmer Zu Wolfsthurn, S. (in press). Dissociable effects of prediction and integration during language comprehension: Evidence from a large-scale study using brain potentials. Philos Trans Royal Soc B.

Nieuwland, M. S., & Van Berkum, J. J. A. (2006). When peanuts fall in love: N400 evidence for the power of discourse. Journal of Cognitive Neuroscience, 18(7), 1098–1111. doi:10.1162/Jocn.2006.18.7.1098

Oldfield, R. C. (1971). The assessment and analysis of handedness: the Edinburgh inventory. Neuropsychologia, 9(1), 97–113. doi:10.1016/0028-3932(71)90067-4

Osterhout, L., & Holcomb, P. J. (1992). Event-related brain potentials elicited by syntactic anomaly. Journal of Memory and Language, 31(6), 785–806. doi:10.1016/0749-596x(92)90039-Z

Osterhout, L., Kim, A., & Kuperberg, G. R. (2012). The neurobiology of sentence comprehension. In M. Spivey, M. Joannisse, & K. McRae (Eds.), The Cambridge Handbook of Psycholinguistics (pp. 365–389). Cambridge: Cambridge University Press.

Osterhout, L., & Nicol, J. (1999). On the distinctiveness, independence and time course of the brain responses to syntactic and semantic anomalies. Language and Cognitive Processes, 14, 283–317.

Paczynski, M., & Kuperberg, G. R. (2011). Electrophysiological evidence for use of the animacy hierarchy, but not thematic role assignment, during verb argument processing. Language and Cognitive Processes, 26(9), 1402–1456. doi:10.1080/01690965.2011.580143

Paczynski, M., & Kuperberg, G. R. (2012). Multiple influences of semantic memory on sentence processing: Distinct effects of semantic relatedness on violations of real-world event/state knowledge and animacy selection restrictions. Journal of Memory and Language, 67(4), 426–448. doi:10.1016/j.jml.2012.07.003

Peirce, J. W. (2007). PsychoPy--Psychophysics software in Python. J Neurosci Methods, 162(1-2), 8–13. doi:10.1016/j.jneumeth.2006.11.017

Qian, T., Jaeger, T. F., & Aslin, R. N. (2012). Learning to represent a multi-context environment: more than detecting changes. Front Psychol, 3, 228. doi:10.3389/fpsyg.2012.00228

Qian, T., Jaeger, T. F., & Aslin, R. N. (2016). Incremental implicit learning of bundles of statistical patterns. Cognition, 157, 156–173.

Quante, L., Bolte, J., & Zwitserlood, P. (2018). Dissociating predictability, plausibility and possibility of sentence continuations in reading: evidence from late-positivity ERPs. PeerJ, 6, e5717. doi:10.7717/peerj.5717

R Core Team. (2016). R: A language and environment for statistical computing. Vienna, Austria: ISBN 3-900051-07-0.

Rabovsky, M., Hansen, S. S., & McClelland, J. L. (2018). Modelling the N400 brain potential as change in a probabilistic representation of meaning. Nature Human Behaviour, 2(9), 693–705. doi:10.1038/s41562-018-0406-4

Rao, R. P., & Ballard, D. H. (1997). Dynamic model of visual recognition predicts neural response properties in the visual cortex. Neural Computation, 9(4), 721–763. doi:10.1162/neco.1997.9.4.721

Rao, R. P., & Ballard, D. H. (1999). Predictive coding in the visual cortex: a functional interpretation of some extra-classical receptive-field effects. Nature Neuroscience, 2(1), 79–87. doi:10.1038/4580

Sanford, A. J., Leuthold, H., Bohan, J., & Sanford, A. J. S. (2011). Anomalies at the borderline of awareness: an ERP study. J Cogn Neurosci, 23, 514–523.

Sassenhagen, J., & Fiebach, C. J. (2019). Finding the P3 in the P600: Decoding shared neural mechanisms of responses to syntactic violations and oddball targets. NeuroImage, 200, 425–436. doi:10.1016/j.neuroimage.2019.06.048

Sassenhagen, J., Schlesewsky, M., & Bornkessel-Schlesewsky, I. (2014). The P600-as-P3 hypothesis revisited: single-trial analyses reveal that the late EEG positivity following linguistically deviant material is reaction time aligned. Brain and Language, 137, 29–39. doi:10.1016/j.bandl.2014.07.010

Schultz, W., & Dickinson, A. (2000). Neuronal coding of prediction errors. Annual Review of Neuroscience, 23, 473–500. doi:10.1146/annurev.neuro.23.1.473

Schumacher, P. B. (2013). When combinatorial processing results in reconceptualization: toward a new approach of compositionality. Front Psychol, 4, 677. doi:10.3389/fpsyg.2013.00677

Schwanenflugel, P. J., & Lacount, K. L. (1988). Semantic relatedness and the scope of facilitation for upcoming words in sentences. J Exp Psychol Learn Mem Cognit, 14(2), 344–354. doi:10.1037//0278-7393.14.2.344

Shetreet, E., Alexander, E. J., Romoli, J., Chierchia, G., & Kuperberg, G. R. (2019). What we know about knowing: Presuppositions generated by factive verbs influence downstream neural processing. Cognition, 184, 96–106. doi:10.1016/j.cognition.2018.11.012

Spratling, M. W. (2008). Predictive coding as a model of biased competition in visual attention. Vision Research, 48(12), 1391–1408. doi:10.1016/j.visres.2008.03.009

Szewczyk, J. M., & Schriefers, H. (2013). Prediction in language comprehension beyond specific words: An ERP study on sentence comprehension in Polish. Journal of Memory and Language, 68(4), 297–314.

Taylor, W. (1953). ’Cloze’ procedure: A new tool for measuring readability. Journalism Quarterly, 30, 415–433.

Thornhill, D. E., & Van Petten, C. (2012). Lexical versus conceptual anticipation during sentence processing: frontal positivity and N400 ERP components. International Journal of Psychophysiology, 83(3), 382–392. doi:10.1016/j.ijpsycho.2011.12.007

Van Berkum, J. J. A., Zwitserlood, P., Hagoort, P., & Brown, C. M. (2003). When and how do listeners relate a sentence to the wider discourse? Evidence from the N400 effect. Cognitive Brain Research, 17(3), 701–718.

van de Meerendonk, N., Kolk, H. H. J., Chwilla, D. J., & Vissers, C. T. W. M. (2009). Monitoring in language perception. Language and Linguistics Compass, 3(5), 1211–1224. doi:10.1111/j.1749-818X.2009.00163.x

van de Meerendonk, N., Kolk, H. H. J., Vissers, C. T. W. M., & Chwilla, D. J. (2010). Monitoring language perception: mild and strong conflicts elicit different ERP patterns. J Cogn Neurosci, 22(1), 67–82. doi:10.1162/jocn.2008.21170

van Dijk, T. A., & Kintsch, W. (1983). Strategies of Discourse Comprehension. New York: Academic Press.

Van Petten, C., & Luka, B. J. (2012). Prediction during language comprehension: benefits, costs, and ERP components. International Journal of Psychophysiology, 83(2), 176–190. doi:10.1016/j.ijpsycho.2011.09.015

Wang, L., Jensen, O., & Kuperberg, G. R. (2018). Neural evidence for prediction of animacy features by verbs during language comprehension: Evidence from MEG and EEG Representational Similarity Analysis. Presented at the 10th Annual Meeting of the Society for the Neurobiology of Language, Quebec City, Canada.

Wicha, N. Y., Moreno, E. M., & Kutas, M. (2004). Anticipating words and their gender: An event-related brain potential study of semantic integration, gender expectancy, and gender agreement in Spanish sentence reading. Journal of Cognitive Neuroscience, 16(7), 1272–1288. doi:10.1162/0898929041920487

Xiang, M., & Kuperberg, G. (2015). Reversing expectations during discourse comprehension. Language, Cognition and Neuroscience, 30(6), 648–672. doi:10.1080/23273798.2014.995679

Yu, A. J., & Dayan, P. (2005). Uncertainty, neuromodulation, and attention. Neuron, 46(4), 681–692. doi:10.1016/j.neuron.2005.04.026

Zirnstein, M., van Hell, J. G., & Kroll, J. F. (2018). Cognitive control ability mediates prediction costs in monolinguals and bilinguals. Cognition, 176, 87–106. doi:10.1016/j.cognition.2018.03.001

Zwaan, R. A., & Radvansky, G. A. (1998). Situation models in language comprehension and memory. Psychological Bulletin, 123(2), 162–185. doi:10.1037/0033-2909.123.2.162

