## Supplementary Materials for "A Tale of Two Positivities and the N400: Distinct neural signatures are evoked by confirmed and violated predictions at different levels of representation"

**Table of Contents**


| | |
|---|---|
| **Table S1.** Lexical characteristics of critical words used in the *high constraint expected* scenarios and the four types of unpredictable scenarios | p. 2 |
| **Table S2.** Pairwise contrasts: N400 (300-500ms, central region) | p. 2 |
| **Table S3.** Pairwise contrasts: *Late frontal positivity* (600-1000ms, prefrontal region) | p. 3 |
| **Table S4.** Pairwise contrasts: *Late posterior positivity/P600* (600-1000ms, posterior region) | p. 3 |
| **Figure S1.** Voltage maps showing N400 effects | p. 4 |
| **Figure S2.** Grand-averaged ERP waveforms at all electrode sites in each of the five conditions | p. 5 |
| **References** | p. 6 |

**Table S1. Lexical characteristics of critical words used in the *high constraint expected* scenarios and the four types of unpredictable scenarios.** The same critical words were counterbalanced across the four types of unpredictable scenarios – *low constraint unexpected*, *high constraint unexpected*, *low constraint anomalous* and *high constraint anomalous*. Different critical words were used in the *high constraint expected* scenarios. Values were retrieved from the English Lexicon Project (Balota et al., 2007), except for Concreteness, which was retrieved from Brysbaert et al. (2014). Log per million frequency values were taken from the SUBTLEX database (Brysbaert & New, 2009). Mean values are shown with standard deviations in parentheses.

| | Number of letters | Log per million frequency | Orthographic Levenshtein Distance | Concreteness |
|---|---|---|---|---|
| Critical words in the four types of unpredictable scenarios | 7.5 (2.23) | 0.6 (0.88) | 2.6 (0.86) | 4.1 (0.69) |
| Critical words in the *high constraint expected* scenarios | 5.7 (1.61) | 1.5 (0.66) | 1.9 (0.56) | 4.3 (0.69) |
| Pair-wise comparison | t(198) = -6.42, p<0.0001 | t(198) = 8.3, p<0.0001 | t(198) = -6.56, p<0.0001 | t(198) = 1.53, p=0.126 |

**Table S2.** Pairwise contrasts: N400 (300-500ms, central region)

| | *High constraint expected* | *High constraint unexpected* | *Low constraint unexpected* | *High constraint anomalous* |
|---|---|---|---|---|
| *High constraint expected* | | | | |
| *High constraint unexpected* | ***t* = 6.06**<br>***p* < .001** | | | |
| *Low constraint unexpected* | ***t* = 6.78**<br>***p* < .001** | *t* = 0.54<br>*p* = .59 | | |
| *High constraint anomalous* | ***t* = 8.47**<br>***p* < .001** | ***t* = 2.07**<br>***p* = .04** | ***t* = 2.74**<br>***p* = .007** | |
| *Low constraint anomalous* | ***t* = 8.99**<br>***p* < .001** | *t* = 1.80<br>*p* = .08 | **t = 2.40**<br>***p* = .02** | t = 0.18<br>*p* = .86 |

All models were run in lmer 4 (1.1-17) using contrast coding, and p-values were obtained using lmerTest (3.0-1) using the Satterthwaite method. All models were maximal (by-subjects and by-items random intercepts and slopes) and there were no convergence errors.

**Table S3.** Pairwise contrasts: *Late frontal positivity* (600-1000ms, prefrontal region)

| | *High constraint expected* | *High constraint unexpected* | *Low constraint unexpected* | *High constraint anomalous* |
|---|---|---|---|---|
| *High constraint expected* | | | | |
| *High constraint unexpected* | **$t$ = 4.10** **$p$ < .001** | | | |
| *Low constraint unexpected* | $t$ = 1.63 $p$ = .11 | **$t$ = 2.37** **$p$ = .02** | | |
| *High constraint anomalous* | $t$ = 0.71 $p$ = .48 | **$t$ = 2.78** **$p$ = .008** | $t$ = 0.68 $p$ = .50 | |
| *Low constraint anomalous* | $t$ = 0.75 $p$ = .45 | **$t$ = 3.37** **$p$ = .001** | $t$ = 0.82 $p$ = .41 | $t$ = 0.2 $p$ = .99 |

**Table S4.** Pairwise contrasts: *Late posterior positivity/P600* (600-1000ms, posterior region)

| | *High constraint expected* | *High constraint unexpected* | *Low constraint unexpected* | *High constraint anomalous* |
|---|---|---|---|---|
| *High constraint expected* | | | | |
| *High constraint unexpected* | $t$ = 1.69 $p$ = .09 | | | |
| *Low constraint unexpected* | $t$ = 1.82 $p$ = .07 | $t$ = 0.21 $p$ = .84 | | |
| *High constraint anomalous* | **$t$ = 6.06** **$p$ < .001** | **$t$ = 4.79** **$p$ < .001** | **$t$ = 5.37** **$p$ < .001** | |
| *Low constraint anomalous* | **$t$ = 5.23** **$p$ < .001** | **$t$ = 3.76** **$p$ < .001** | **$t$ = 3.85** **$p$ < .001** | **$t$ = 2.31** **$p$ = .03** |

All models were run in lmer 4 (1.1-17) using contrast coding, and p-values were obtained using lmerTest (3.0-1) using the Satterthwaite method. All models were maximal (by-subjects and by-items random intercepts and slopes) and there were no convergence errors.

**Figure S1. Voltage maps showing N400 effects**. Voltages are averaged across the 300-500ms time window for all pair-wise contrasts.

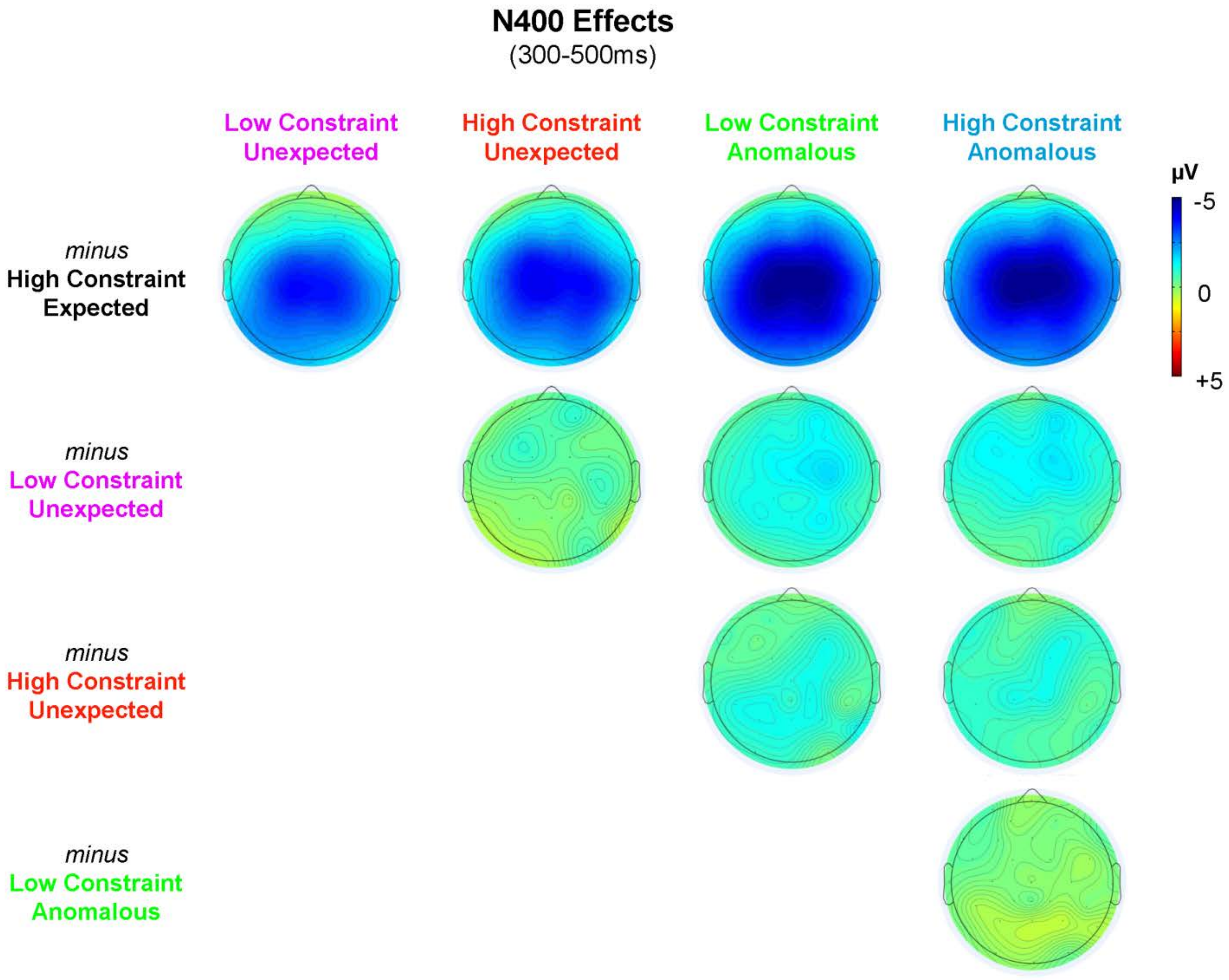

**Figure S2.** Grand-averaged ERP waveforms at all electrode sites in each of the five conditions. Negative voltage is plotted upward.

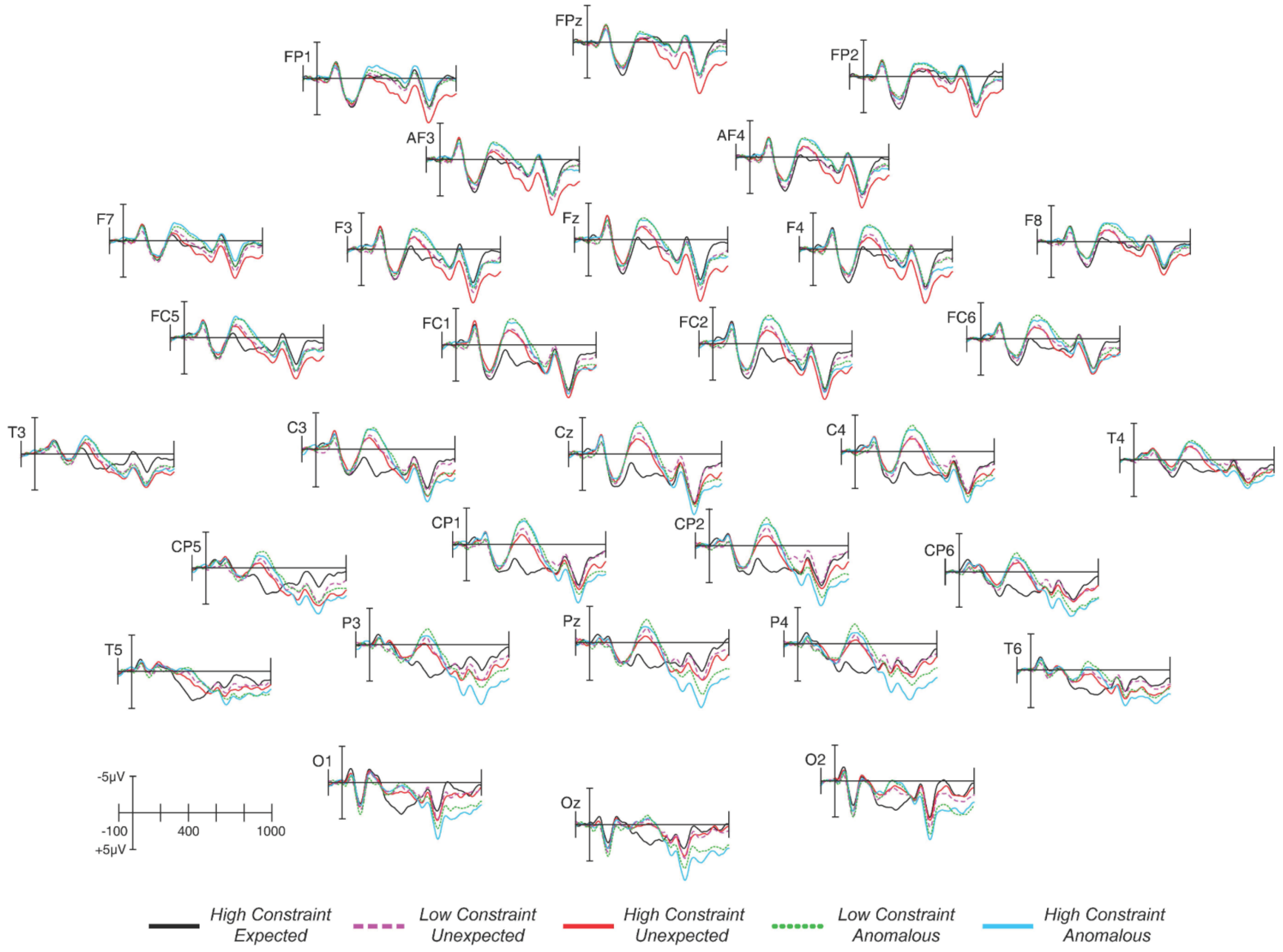

### References

Balota, D. A., Yap, M. J., Hutchison, K. A., Cortese, M. J., Kessler, B., Loftis, B., . . . Treiman, R. (2007). The English lexicon project. *Behav Res Methods, 39*(3), 445-459.

Brysbaert, M., & New, B. (2009). Moving beyond Kučera and Francis: A critical evaluation of current word frequency norms and the introduction of a new and improved word frequency measure for American English. *Behavioral Research Methods, 41*, 977-990.

Brysbaert, M., Warriner, A. B., & Kuperman, V. (2014). Concreteness ratings for 40 thousand generally known English word lemmas. *Behav Res Methods*, *46*(3), 904-911. doi:10.3758/s13428-013-0403-5
